# TET2-mediated epigenetic reprogramming of breast cancer cells impairs lysosome biogenesis

**DOI:** 10.1101/2021.10.27.466063

**Authors:** Audrey Laurent, Thierry Madigou, Maud Bizot, Marion Turpin, Gaëlle Palierne, Elise Mahé, Sarah Guimard, Raphaël Métivier, Stéphane Avner, Christine Le Péron, Gilles Salbert

## Abstract

Methylation and demethylation of cytosines in DNA are believed to act as keystones of cell-specific gene expression through controlling chromatin structure and accessibility to transcription factors. Cancer cells have their own transcriptional programs and we sought to alter such a cancer-specific program by enforcing expression of the catalytic domain (CD) of the methylcytosine dioxygenase TET2 in breast cancer cells. TET2 CD decreased the tumorigenic potential of cancer cells through both activation and repression of a repertoire of genes that, interestingly, differed in part from the one observed upon treatment with the hypomethylating agent decitabine. In addition to promoting the establishment of an antiviral state, TET2 activated 5mC turnover at thousands of MYC binding motifs and down-regulated a panel of known MYC-repressed genes involved in lysosome biogenesis and function. Thus, an extensive cross-talk between TET2 and the oncogenic transcription factor MYC establishes a lysosomal storage disease-like state that contributes to an exacerbated sensitivity to autophagy inducers.

## INTRODUCTION

Cell-specific gene expression programs are sustained by epigenetic landscapes that are established by enzymes targeting histones and DNA. Accordingly, genome-wide epigenomic rewiring associates with acquisition of new cellular identities during development (1). Cancer cells, although maintaining a cell-of-origin epigenomic imprint, acquire specific epigenomic features, some of which are common between different cancer types (2–4). One such cancer-associated epigenetic feature is the so-called CpG Island Methylator Phenotype (CIMP) in which hypermethylation of a substantial number of CpG islands (CGIs) that surround transcription start sites (TSSs) is found to be associated with low gene expression of the corresponding genes (3, 5). In agreement with the idea that CGI methylation can occur at similar positions in various cancers, CGIs from the clustered proto-cadherin (PCDH) tumor-suppressor locus are frequently found methylated in breast, Wilm’s tumor, cervical, colorectal, gastric, and biliary tract cancers (6–9). Although the existence of a breast cancer CIMP (B-CIMP) has been debated, a phenotype comparable to colon cancer and glioma CIMP has been evidenced and suggested to be prevalent in the estrogen receptor *α* (ER*α*) and progesterone receptor (PR) positive luminal subtype of breast tumors (10). Such an association of B-CIMP with the ER*α* and PR status was later confirmed, but no correlation with tumor size, lymph node invasion and metastasis could be evidenced (11). Although it is not precisely known what triggers CIMP, a decrease in the activity of Ten Eleven Translocation (TET) enzymes has been documented in various cancers (12) and linked to the occurrence of CIMP in leukemia (13) and colorectal cancer (14). TETs are 2-oxoglutarate/Fe^2+^-dependent dioxygenases that iteratively oxidize 5-methylcytosine (5mC) into 5-hydroxymethylcytosine (5hmC), 5-formylcytosine (5fC) and 5-carboxylcytosine (5caC), 5fC and 5caC being replaced by unmodified cytosines by the successive action of the DNA glycosylase TDG and the base excision repair machinery (15). Consistent with a role in maintaining an hypomethylated state in CpG-rich regions, TET1 knock-out in mouse embryonic stem cells (mESCs) leads to CGI hypermethylation, suggesting that CIMP could indeed be caused by a reduced TET activity in cancer cells (16). However, TET1 engagement at promoters can also repress gene expression by favoring the recruitment of PRC2, a complex mediating H3K27 methylation (16). This dual action of TET enzymes towards gene regulation suggests that enforcing TET activity in cancer cells could provide additional benefits compared to epigenetic drugs commonly used to inhibit DNA methylation like 5-aza-cytidine and 5-aza-deoxycytidine (decitabine). Here, we genetically engineered MCF-7 cells, a model of low metastatic luminal breast cancer cells that express both ER*α* and PR transcription factors and whose growth is partly dependent on estrogen supply (17), to artificially modify their epigenome. Enforcing TET2 activity reduced the tumorigenic potential of MCF-7 breast cancer cells, and triggered an antiviral state and a lysosomal storage disease-like phenotype that prediposed cells to death.

## MATERIALS and METHODS

### Cell culture and reagents

MCF-7 cells stably transfected with an empty vector or with plasmids encoding either wild type mouse TET2 catalytic domain (CD - aa916-1921, pcDNA3-Flag-TET2 CD, addgene #72219, 18) or the catalytically dead mutant H1304Y, D1306A (pcDNA3-Flag-TET2 mCD, addgene #72220, 18) were grown in high glucose and pyruvate containing DMEM (Gibco 41966) supplemented with 10% fetal calf serum (Eurobio S116365181H), non-essential amino acids (Gibco 11140035), Penicillin-Streptomycin (Gibco 15240) and Geneticin (Gibco 11811064) at 37°C and 5% CO2. The SIRT1 activator compound SRT1720 was purchased from Sigma-Aldrich (567860) and chloroquine was from the CYTO-ID Autophagy Detection kit 2.0 (ENZO ENZ-KIT175). Decitabine (5-aza-deoxycytidine) was from SIGMA-ALDRICH (A3656). For RT-qPCR analysis, cells were treated with 100 nM decitabine given every 36 hours, for 96 hours.

### Cell cycle analysis, migration and clonogenicity assays

Cell cycle was analysed by flow cytometry. Briefly, 2,000,000 cells were plated on 10-cm dishes in DMEM supplemented with 10% FBS. After 72 hours, cells were trypsinized and fixed with 70% ethanol before being stained with propidium iodide in the presence of RNAse A. Cells were acquired on a FORTESSA Beckton Dickinson cytometer (Flow Cytometry Biosit Facility) and cell cycle analysis was performed with BD FACS DIVA software. For EdU (5-Ethynyl-2’-deoxyuridine) labeling of S phase cells, 250,000 cells were plated on coverslips in 6-well plates. After 24 hours the cells were incubated with serum- and steroid-deficient medium and were then treated with 10 nM E_2_ or vehicule for 24 hours in a 0.5% serum-supplemented medium. Incorporated EdU was then fluorescently labeled with Alexa Fluor^TM^ 488 by click chemistry (Click-iT® EdU imaging Kit, Invitrogen) according to the manufacturer instructions. For wound healing assays, 650.10^3^ cells were seeded into 10 mm^2^ dishes. After an overnight culture, cells were starved for 72h. Confluent starved cells were wounded with a pipette tip and treated with E2 (10nM) or ethanol. Images of the recovery were captured daily after the renewal of the medium containing E2 (10nM) or ethanol. For soft agar assays, respectively 5,000, 10,000 or 20,000 cells were seeded on soft agar 10 cm dishes (respectively 0,5 % and 0,33 % agar for down and top layers in complete medium). After 4 weeks of culture (adding twice a week 500 μL of complete medium to avoid desiccation), colonies were stained with 0.005% crystal violet in 2% ethanol, imaged and counted.

### siRNA transfection and cell viability determination

Levels of ABCE1 messenger RNAs (mRNAs) were reduced by reverse transfection with 27-mer duplex siRNAs targeting ABCE1 (OriGene, SR304089B: rArGrArArGrUrArCrCrArGrUrUrCrUrArArArUrGrUrCrAGT, SR304089C: rGrGrCrUrArGrArArArGrUrArUrGrGrUrUrUrArArCrUrGGA). Increasing concentrations (0, 2.5, 5 and 10 nM) of scrambled (scr, OriGene, SR30004) or ABCE1 siRNAs diluted in OptiMEM (Thermofisher, 31985070) were transfected in triplicates in 48-well plates with Lipofectamine RNAiMAX (Thermofisher, 13778075) before seeding EV, TET2 CD and TET2 mCD cells (2.5.10^4^ in each well). After 48 hours, cell viability was quantified with a MTT detection kit (Abcam, ab211091). Medium was replaced by 100 μL of a 1:1 mix of phenol-red free DMEM without serum and a 10 % MTT solution, and the plate was further incubated for 3 hours at 37°C. The formed insoluble formazan cristals were then dissolved with a 1:1 solution of dimethylsulfoxide (DMSO) (Sigma, D8418) and isopropanol (Sigma, 33539). After 30 min of incubation and a transfer into a 96-well plate, absorbance was detected at OD=570nm using a Bio-TEK microplate reader (Power wave XS).

### Detection of acidic vesicles

Acidic vesicles were labeled with LysoTracker Red (ThermoFisher L7528). Cells were seeded on glass coverslips and grown in complete medium (DMEM, 10% serum) for 24 hours before adding Lysotracker Red (ThermoFisher) directly in the medium (final concentration: 75 nM) and Hoechst 33342 for nuclear staining. After 30 minutes at 37°C, cells were washed once with PBS and fixed for 15 minutes with 4% paraformaldehyde in PBS. After two washes with PBS, coverslips were mounted on a glass slide with Vectashield (Vector Laboratories H-1200). Cells were imaged with an Olympus BX-51 fluorescence microscope (60x magnification) and images were processed with Image J for quantification. Briefly, color channels were split and the red channel images were processed with “Find edges” before threshold adjustment and particle size determination. Statistical differences (t test) were determined with Prism 5 (GraphPad Software Inc.).

### Dot blot detection of 5hmC

Genomic DNA (gDNA) was prepared using the DNeasy Blood and Tissue kit (QIAGEN 69506). Relative 5hmC levels were quantified blotting 500 ng of gDNA on a nitrocellulose membrane with an anti-5hmC antibody (Active Motif, 39769) diluted 1/10000 followed by an anti-rabbit antibody coupled to horseradish peroxydase (Dutscher NA934) diluted 1/5000. DNA was stained with 0.04% methylene blue in 0.5 M sodium acetate.

### RNA preparation and dot blot detection of dsRNAs

Total RNAs were extracted from 5.10^7^ EV, TET2 CD and TET2 mCD cells using TRIzol reagent (Thermofisher scientific 15596018) according to the manufacturer’s instructions. Two thousand nanograms, 1000 ng and 500 ng of each RNA sample were spotted to a nylon membrane (Hybond N, Dutscher RPN203B) previously soaked in 2X SSPE solution (0.3M NaCl, 20mM sodium phosphate, 2 mM EDTA) and inserted in the dot blot apparatus (SCIE-PLAS). RNAs were crosslinked to the membrane by 30 min heating at 80°C. Membranes were incubated overnight with an anti dsRNA antibody (dsRNA mAb J2, Scicons) diluted 1/500 followed by an anti-mouse antibody coupled to horseradish peroxidase (Santa Cruz, sc-2005) diluted 1/10000. Total RNA levels were stained with 0.04 % methylene blue in 0.5 M sodium acetate.

### RT-qPCR analyses

Total RNAs were isolated from 5.10^7^ cells using TRIzol*®* reagent (Thermofisher, 15596018) according to the manufacturer’s protocol. Reverse transcription was performed using 500 ng of total RNAs as template, 200 units of M-MLV reverse transcriptase (ThermoFisher 28025013) and 250 ng of Pd(N)6 random hexamers (Euromedex PM-301L). Real-time qPCR of reversed transcribed RNAs was run with SYBR*®* Green Master Mix (Biorad, 1725006CUST) in a Bio-Rad CFX96. Data were normalized to the positive control CDK8 according to the 2^-ΔΔCt^ method. Primers were designed using the software Primer3 (http://frodo.wi.mit.edu/primer3/; (Untergasser et al., 2012)) and were synthetized by SIGMA-ALDRICH.

Primer sequences:

DSCR8_Up CAAAAATGAAGGAGCCTGGA

DSCR8_Do ACCACTGCACTCCAGCAGTA

MCAM_Up CGCACACAGCTGGTCAAC

MCAM_Do CCTGACGCTTCACAAGACAG

SEMA6B_Up GTGCGCCAACTACAGCATAG

SEMA6B_Do AAGAGCATCCCGTCAGAGAA

SOX2_Up TACAGCATGATGCAGGACCA

SOX2_Do GTACTGCAGGGCGCTCAC

ABCE1_Up TTGCAGAGATTTGCTTGTGC

ABCE1_Do GCCTTTAAACGCTGCTTGAC

DDX58 1_Up TGTTGAAACAGAAGATCTTGAGGA

DDX58 1_Do CACTTCTGAAGGTGGACATGAA

OAS1_Up CTGAGGCCTGGCTGAATTAC

OAS1_Do TCGTCTGCACTGTTGCTTTC

OAS2_Up AGAATCTCTTTCGAGGTGCTG

OAS2_Do CCAGGACTGGCATTTGTCTT

IFIT1_Up ACACCTGAAAGGCCAGAATG

IFIT1_Do TCCTCACATTTGCTTGGTTG

CDK8 Up TCCTGCAGATAAAGATTGGGAAG

CDK8 Do CTTGATAAGGCTGCAGTTGGT

CTSD_Up CACAAGTTCACGTCCATCCG

CTSD_Do TGCCAATCTCCCCGTAGTAC

JPH3_Up CCAGAAATCCTTGCCTGTCG

JPH3_Do GCAAGATCACCATGACCACC

LHX2_Up CAAGATCTCGGACCGCTACT

LHX2_Do GAAACAGGTGAGCTCCGACT

H19_Up TCCTGAACACCTTAGGCTGG

H19_Do TCATGTTGTGGGTTCTGGGA

CLN3_Up CGTCCTGGTTGCCTTTTCTC

CLN3_Do CAGTGAGGGAGAGGAAGGTG

NAGLU_Up GAGATAGACTGGATGGCGCT

NAGLU_Do TTGATCTCTGCCTGGGTCAG

HERV-FC1_Up TTTCCCACCGCTGGTAATAG

HERV-FC1_Do AGGCTAAGGATTCGGCTGAG

LTR12C_Up AGCTAGACATAAAGGTCCTCCACG

LTR12C_Do TGGGGCCTTGGAGAACTTTTATG

### ChIP-seq

All experiments were performed under hormonal depletion: cells were kept for 48h in phenol-red-free DMEM (Gibco 31052) supplemented with 2.5% dextran-charcoal-treated fetal calf serum (Eurobio S116365181W), glutamine, sodium pyruvate, non-essential amino acids, Penicillin-Streptomycin and Geneticin at 37°C and 5% CO2. For H3K4me3 and H3K27me3 ChIP-seq experiments, 4 million cells were fixed in 1.5% formaldehyde (Sigma F8775) for 10 min at room temperature, the reaction was stopped by the addition of glycine (100 mM). Cells were lysed in lysis buffer (150 mM Tris-HCl pH 8.1, 10 mM EDTA, 0.5% Empigen BB, 1% SDS, protease inhibitor cocktail (Roche 5056489) and sonicated using a bioruptor (Diagenode, 15 min 30 sec ON /30 sec OFF). Sonicated chromatin was incubated at 4°C overnight with either an Anti-H3K4me3 (Millipore 04-745; 1 µg) or Anti-H3K27me3 (Millipore 07-449; 1 µg) antibody in IP buffer (2.8 ng/mL yeast tRNA, 20 mM Tris-HCl, 2 mM EDTA, 150 mM NaCl, 1% Triton X-100, proteases inhibitor cocktail). Complexes were recovered after incubation with 50 µl protein A-conjugated sepharose beads slurry at 4°C. Beads were washed in washing buffers WB1 (20 mM Tris-HCl pH 8, 2 mM EDTA, 150 mM NaCl, 0.1% SDS, 1 % Triton X-100), WB2 (20 mM Tris-HCl pH 8, 2 mM EDTA, 500 mM NaCl, 0.1% SDS, 1 % Triton X-100), WB3 (10 mM Tris-HCl pH 8, 1 mM EDTA, 250 mM LiCl, 1 % deoxycholate, 1 % NP-40), WB4 (10 mM Tris-HCl pH 8, 1mM EDTA) and fragments were eluted in extraction buffer (1% SDS, 0.1 M NaHCO3). For the preparation of each sequencing library (TruSeq, Illumina), ChIPed DNA from 9 independent ChIP experiments were pooled and sequenced at the GenomEast Platform (IGBMC, Strasbourg). Primers used for H3K27me3 ChIP-qPCR were synthetized by SIGMA-ALDRICH. Primer sequences were:

JPH3_forward GAGTCCGTTTTCACCGTTTG

JPH3_reverse ACCCTCCGTCGTCAAAATTA

LHX2_forward GCGCACTGATCAATCACC

LHX2_reverse GGAGAAAGTGAGGCCAGACC

### RNA-seq

RNA-seq was performed in triplicates on single EV, TET2 CD, and TET2 mCD clones after 4 h of treatment with E_2_ (10 nM)/ethanol of cells previously maintained in phenol-red free medium supplemented with 2.5% charcoal-treated fetal calf serum during 48 hrs. Total RNA extraction was run using RNeasy Plus kit (QIAGEN) which includes an optional DNAse digestion. For each condition, three replicate libraries (TruSeq stranded mRNA) were prepared and sequenced (single reads of 75 bases) at the Genomic Paris Center facility (Paris, France). For RNA-seq statistical analysis, one replicate of the TET2-mCD control RNA-seq was ignored due to its lack of similarity with the other two samples of the triplicate, according to PCA analysis and hirerachical clustering.

### SCL-exo-seq

Selective Chemical Labelling-exonuclease (SCL-exo, 19) experiments were conducted on starved cells after 50 min of E2 (10nM)/ethanol treatment. Genomic DNA was extracted using the DNeasy Blood and Tissue kit (QIAGEN 69506). For each experiment, 8 μg of gDNA were sonicated two times 7 min (30 sec ON /30 sec OFF) and two times 14 min (30 sec ON /30 sec OFF) with a bioruptor (Diagenode). Glucosylation and biotinylation of 5hmC were performed with the Hydroxymethyl Collector kit (Active Motif 55013), followed by on beads-exonuclease digestion of the captured fragments and library preparation (TrueSeq library preparation kit, Ilumina IP-202-1012). For normalization purpose, 400 pg of 5hmC Control DNA provided by the Hydroxymethyl Collector kit were added to each sample as spike-in. Libraries from 7 independent SCL-exo experiments were sequenced on 7 lanes of a HiSeq 1500 (Illumina) by the GEH facility (Rennes, France).

### Quantification and statistical analyses

RNA-seq reads were mapped to hg19 with Bowtie (20) and transcripts were quantified with RSEM (21). Differentially expressed genes were identified from RNA-seq data by the R package DESeq2 (22) after filtering of the raw data using the R package HTSFilter (23). Online tools (GREAT, 24; http://bejerano.stanford.edu/great/public/html/; Panther, 25; http://www.pantherdb.org/) were used for interpretation and functional annotation. ChIP-seq reads were mapped to hg19 using Bowtie (20). SAMtools (26) generated bam files which were processed with MACS (27) to generate wig files. Peak calling followed a previously described procedure (28). Sequencing reads from publicly available datasets were mapped and treated following the same procedure as above. MCF-7 MeDIP sequencing reads (DRA000030, 29) and MYC ChIP-seq reads (SRR575112) were downloaded from http://trace.ddbj.nig.ac.jp/dra/index_e.shtml and from https://www.ncbi.nlm.nih.gov/sra?term=SRX188954, respectively. Differential wig files were generated by substracting the signal in EV cells from the signal in either TET2 CD or TET2 mCD cells after normalization to the number of reads. ER*α* ChIP-seq heatmaps were generated online with Cistrome (30; http://cistrome.org/ap/root). SCL-exo reads were mapped separatly on both strands with Bowtie (20). The resulting SAM files were processed with a python script (https://mycore.core-cloud.net/index.php/s/4gyZ9dLTqgo86dt) to identify 5hmCpGs (31). ChIP-seq and SCL-exo data were normalized to the input as follows: for every position of the wig file, a window of 100 bp surrounding that position was considered and the input signal values in that window were averaged. Sample signal values were next divided by the mean input value of their corresponding window. For transcription factor motif search, bed files containing the coordinates of CpGs identified by SCL-exo were analyzed online with TFmotifView (32). The Cancer Genome Atlas (TCGA) RNA-seq data from breast cancer patients were downloaded from UCSC Xena (https://xenabrowser.net/). Heatmaps shown in Supplementary Fig. 5 were generated online by UCSC Xena (https://xenabrowser.net/heatmap/). RNA-seq data from intrinsic molecular breast cancer subtypes (PAM50) were interrogated with bc-GenExMiner v4.6 (33; http://bcgenex.centregauducheau.fr/BC-GEM/GEM-Accueil.php?js=1). Violin plots from Fig. 1, Fig. 2 and Supplementary Fig. 1 were generated online by bc-GenExMiner v4.6. Statiscal differences were analysed with a Dunnett-Tukey-Kramer’s test. Venn diagrams comparing gene lists were generated online (http://bioinformatics.psb.ugent.be/webtools/Venn/). Bar graphs were generated with GraphPad Prism 5.0 and analyzed by unpaired t-test (Prism 5.0). In each case, number of samples are indicated in the corresponding figure legend.

**Figure 1:**
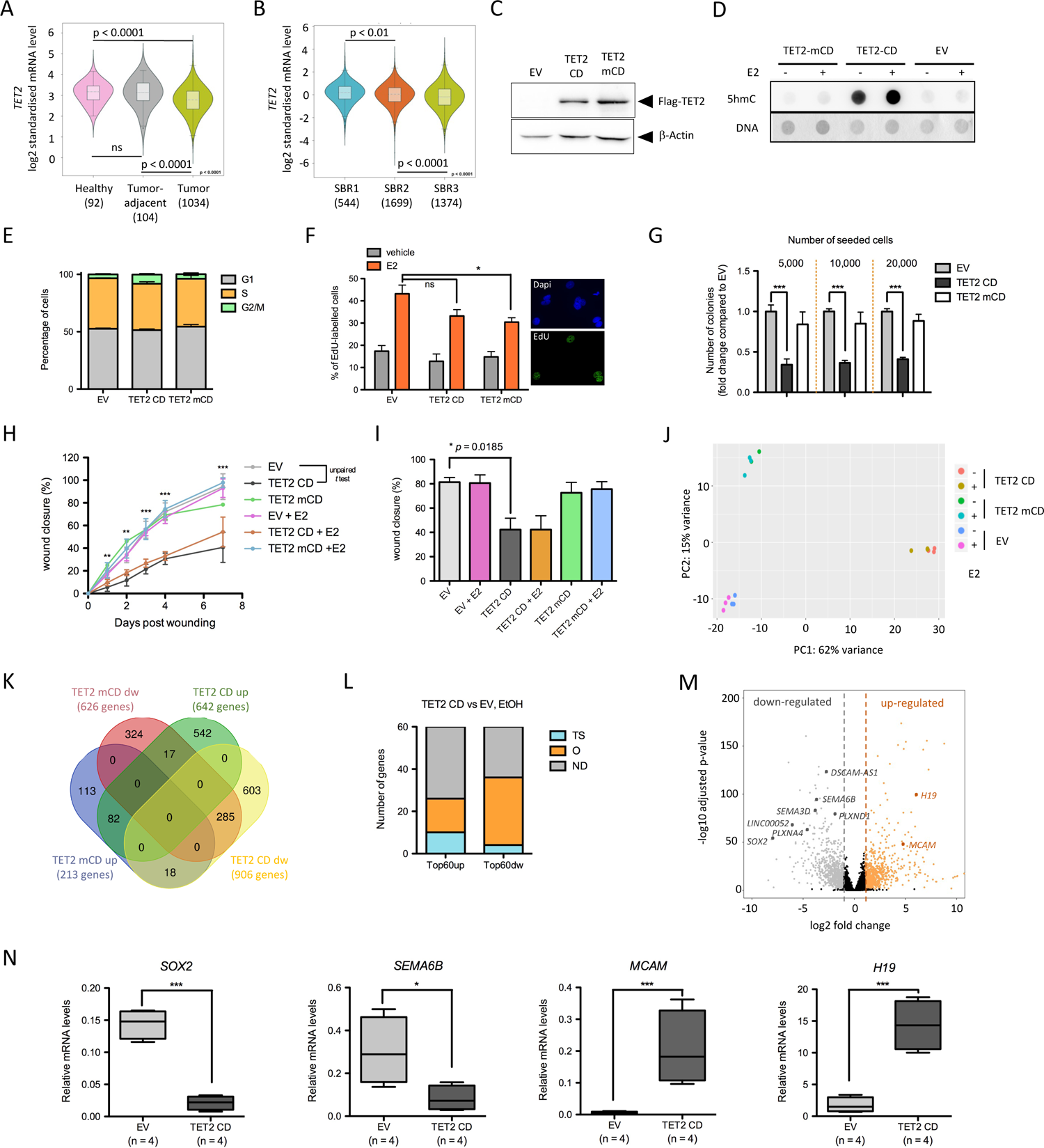
TET2 CD expression reduces the tumorigenic potential of MCF-7 cells. (**A**) TET2 expression in healthy tissue, tumor adjacent and tumors of BRCA patients. (**B**) TET2 expression in BRCA tumors according to their Scarff Bloom and Richardson grade status. In A and B, plots were generated with Breast Cancer Gene-Expression Miner v4.5. (**C**) Western blot detection of Flag-TET2 CD and TET2 mCD in MCF-7 clones. (**D**) Dot blot analysis of 5hmC levels in 500 ng of genomic DNA from EV, TET2 CD and TET2 mCD cells treated or not with estradiol (E2). DNA was stained with methylene blue. (**E**) Flow cytometry of propidium iodide-labeled EV, TET2 CD and TET2 mCD cells. Bar graphs show the distribution of cells in G1, S and G2/M phases of the cell cycle (n=5). (**F**) Bar graph representation of EdU labeling of S phase cells in the presence or absence of E2 (n=3). Images show dapi staining and EdU labeling of a representative microscopic field of EV cells treated with E2. (**G**) Anchorage-independent clonogenicity assay of MCF-7-derived clones grown in soft agar for 4 weeks. Bar graphs indicate the number of colonies (mean +/- SEM, n=6) for three initial seeding densities. (**H,I**) Wound-healing assay showing delayed migration of TET2 CD cells. Wound closure was quantified at different time points after wounding and data are shown as mean +/- SEM (n=6) for all time points from a single experiment (**H**) or as mean +/- SEM (n=3) for day 4 samples from three independent experiments (**I**). (**J**) Principal component analysis of RNA-seq samples based on the 500 most expressed genes in each sample. (**K**) Venn diagram showing the overlap between the lists of up- or down (dw)-regulated genes in TET2 CD and TET2 mCD cells compared to EV cells in the absence of E2 (FC ≥ 2). (**L**) Literature-based annotation of the top 60 up- and down-regulated genes without E2 in TET2 CD cells compared to EV cells (TS: tumor suppressor, O: oncogene, ND: not determined). (**M**) Volcano plot visualization of differentially expressed genes (DEGs) between TET2 CD and EV cells. Genes described in the text have been highlighted. Dashed vertical lines indicate −1 and +1 log2 fold change. (**N**) Box plot representation of RT-qPCR measurements of *SOX2*, *SEMA6B*, *MCAM* and *H19* RNAs in 4 independent EV and TET2 CD clones (mean +/- SEM, n=4).

**Figure 2:**
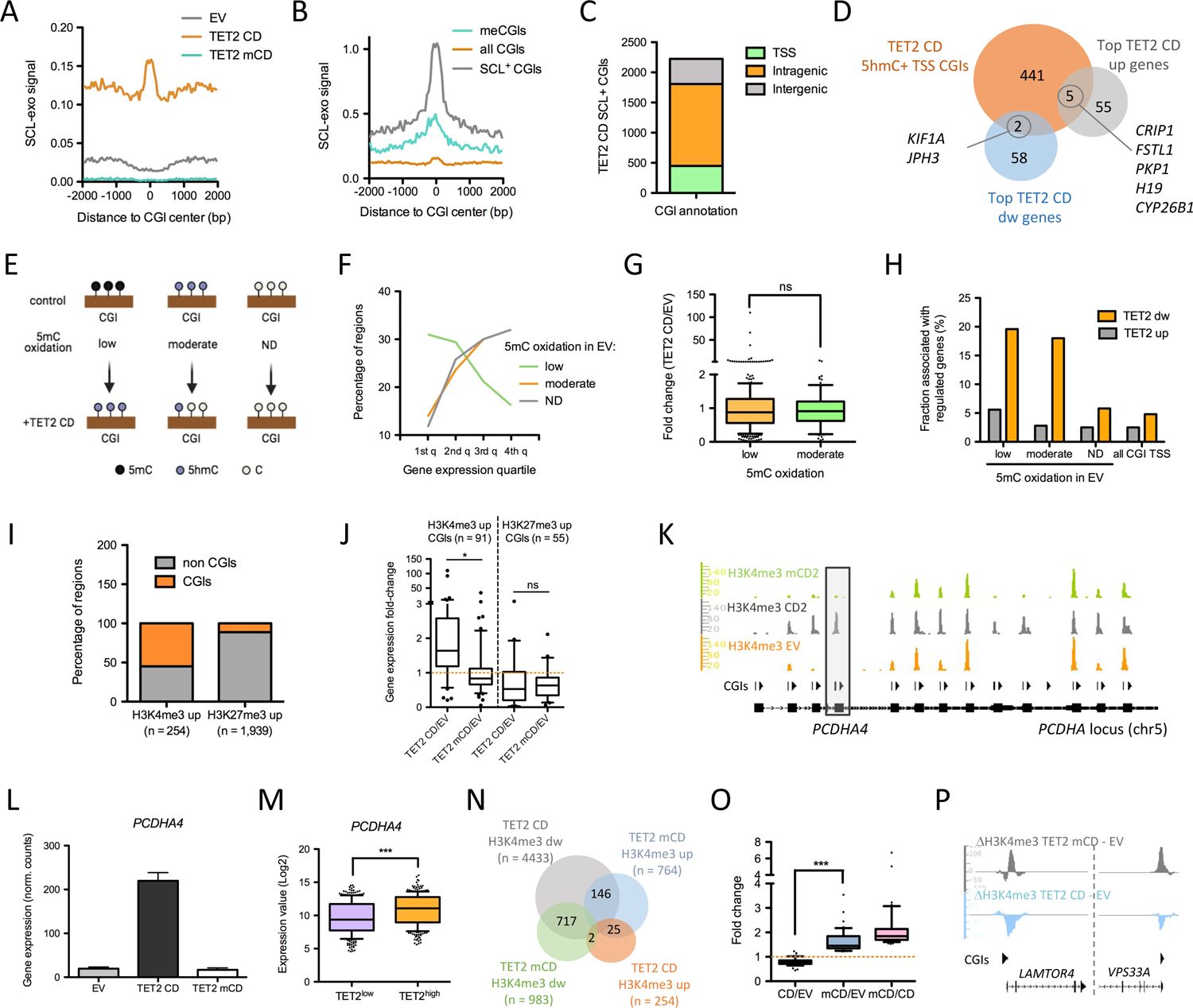
CGI reprogramming by TET2 CD. (**A**) Average profile of 5hmC levels (SCL-exo signal) in EV, TET2 CD and TET2 mCD cells, centered at hg19 CGIs (n=28,691). (**B**) Average profile of 5hmC levels in TET2 CD cells centered at all hg19 CGIs, at CGIs methylated in MCF-7 cells (n=2,330) or at SCL-positive CGIs in TET2 CD cells (n=2,224). (**C**) Called 5hmC positive CGIs in TET2 CD cells were annotated as intergenic, intragenic or within +/- 500 bp of a TSS. (**D**) Venn diagram visualization of the overlap between Top 60 DEGs (up and down) and 5hmC-positive TSS-associated CGIs in TET2 CD cells. (**E**) Classification of TSS-CGIs as a function of 5hmC variation between EV and TET2 CD cells. Low 5mC oxidation TSS-CGIs have no detectable 5hmC in EV cells and a gain in TET2 CD cells. Moderate 5mC oxidation TSS-CGIs have detectable 5hmC signal in EV an a decreased signal in TET2 CD cells. TSS-CGIs with no detectable (ND) 5hmC either in EV cells or in TET2 CD cells are supposed to be protected from DNA methylation. (**F**) Association of the classified TSS-CGIs with gene expression quartiles (1st quartile including lowest expression levels to the 4th quartile with highest expression levels). (**G**) Expression fold change (TET2 CD/EV) of genes associated with low or moderate 5mC oxidation TSS-CGIs. (**H**) Fraction of low, moderate and ND 5mC oxidation TSS-CGIs associated with up- and down-regulated genes in TET2 CD cells compared to EV cells. (**I**) CGI association of H3K4me3 up or H3K27me3 up regions in TET2 CD cells *versus* EV cells. (**J**) Expression fold change of gene associated with H3K4me3 up or H3K27me3 up CGIs in TET2 CD or TET2 mCD versus EV cells. (**K**) IGB visualization of H3K4me3 ChIP-seq signal at the *PCDHA* locus in EV and TET2 CD and TET2 mCD cells. CGI positions are indicated. (**L**) *PCDHA4* gene expression levels in EV, TET2 CD and mCD cells (normalized RNA-seq read counts, mean +/- SEM, n=3). (**M**) *PCDHA4* mRNA levels in breast cancer tumours (TCGA BRCA panel) classified into TET2^low^ (1st quartile n=301) and TET2^high^ (4th quartile, n=299; samples with a value of 0 were discarded). (**N**) Venn diagram indicating the overlap between regions that gained (up) or lost (dw) H3K4me3 in TET2 CD and mCD cells compared to EV cells. (**O**) Expression fold change of the 42 genes showing opposite H3K4me3 signal variations in TET2 CD and mCD cells compared to EV cells and that were up-regulated more than 1.2 fold in TET2 mCD versus EV cells and more than 1.5 fold in TET2 mCD versus TET2 CD. (**P**) IGB visualization of the differential H3K4me3 signal at the *LAMTOR4* and *VPS33A* loci.

## RESULTS

### Active TET2 CD alters the tumorigenic potential of MCF-7 cells

Examination of TET gene expression in breast cancer patients revealed that although TET3 mRNA levels were higher in tumors and positively correlated with tumor progression, both TET1 and TET2 expression were diminished in tumor cells, and TET2 expression decreased with tumor progression more significantly than TET1 (Fig. 1A,B and Supplementary Fig. 1A). We thus chose to ectopically express TET2 in breast cancer cells and transfected MCF-7 cells with an expression vector for the Flag-tagged active murine TET2 CD and, as controls, with a catalytically dead mutant (H1304Y and D1306A) TET2 mCD (18) or an empty vector (EV). Clones were isolated in the presence of Geneticin and analyzed for expression of Flag-TET2 (Fig. 1C) as well as for their enrichment in 5hmC (Fig. 1D). Consistent with a role of TET2 in 5mC oxidation, increased 5hmC levels were observed in TET2 CD cells but not in TET2 mCD cells. Although TETs have been suggested to regulate cell cycle (34), no significant changes in the distribution of the cells in the different cell cycle phases were noticed by flow cytometry analysis (Fig. 1E). In addition, S phase entry was still enhanced by estradiol in TET2 CD and mCD cells, as assessed by EdU labelling (Fig. 1F). Conversely, TET2 CD cells were less prone to grow as colonies in an anchorange-independent growth assay (Fig. 1G), and to migrate in a wound healing assay (Fig. 1H,I). Collectively, these data show that enforcing active TET2 CD expression mitigates the tumorigenicity of MCF-7 cells and are consistent with previous work showing that both TET1 and TET2 reduce tumor growth in xenograft mice (35, 36).

To explore further the impact of TET2 CD and mCD expression in the MCF-7 clones, their transcriptome was established through Illumina sequencing of poly-dT-captured mRNAs. Principal component analysis (PCA) of the 500 most expressed genes indicated a major reconfiguration of RNA pol II-mediated transcription in TET2 CD cells and to a lower extent in TET2 mCD cells, with little impact of estradiol (Fig. 1J). Comparison of TET2 CD and mCD differentially expressed genes (DEGs, fold change [FC] ≥ 2 and adjusted *p* value ≤ 0.05 when compared to EV) evidenced that the catalytic activity of TET2 was not required for all of the transcriptional changes observed (Fig. 1K). Indeed, although the number of regulated genes was lower in the case of TET2 mCD, 12.7% (82 genes) of the 642 genes activated by TET2 CD were also activated by expression of TET2 mCD and 31.4% (285 genes) of the 906 genes repressed by TET2 CD were also repressed by TET2 mCD. This suggests that TET2 CD can exert 5mC oxidation-independent functions that are more prominent for gene repression than for gene activation but also indicates that 5mC oxidation *per se* is an important determinant of TET2 CD-mediated gene regulation. Close examination of the top 60 differentially regulated genes in TET2 CD cells *versus* EV cells revealed that a rather similar proportion of described oncogenes (26.7%), including the long non-coding RNA (lncRNA) *H19* (37 - Fig. 1L,M,N) and tumor suppressor genes (16.7%) were up-regulated (Fig. 1L and Supplementary Table 1). Conversely, 56.7% of the top 60 down-regulated genes were known oncogenes (Fig. 1L and Supplementary Table 1), among which the lncRNAs *DSCAM-AS1*, a luminal marker involved in breast cancer progression (38, 39), and *LINC00052*, an anchorage-independent growth promoter (40 - Fig. 1M). Noticeably, 4 genes encoding semaphorins (SEMA) of the oncongenic type and their receptors (PLXN) were among the top 60 down-regulated genes in TET2 CD cells (Fig. 1M). *SEMA3D* and *SEMA6B* as well as the receptors *PLXNA4* and *PLXND1* act as oncogenes favoring cell growth and migration (41–43). Interestingly, the mixed oncogene/tumor suppressor *MCAM*, which is induced through interaction of the tumor suppressor type semaphorin SEMA3A with its receptor NPR1 and silenced by promoter methylation in MCF-7 cells (44, 45), was among the top 60 TET2 CD up-regulated genes (Fig. 1M). In addition, a dramatic reduction in expression of the major oncogene *SOX2* (46, 47) was evidenced (Fig. 1M). Additional TET2 CD clones were tested by RT-qPCR and consistently showed *SOX2* and *SEMA6B* down-regulation, and up-regulation of *MCAM* and *H19* (Fig. 1N). Except for *SOX2*, gene expression changes for *SEMA6B*, *MCAM*, and *H19* in MCF-7 cells treated with the hypomethylating agent decitabine (48) were similar to those observed upon TET2 CD expression, validating the hypothesis that these genes are regulated by DNA methylation (Supplementary Fig. 1B). Finally, consistent with a lower tumorigenicity of TET2 CD cells, interrogation of proteomic data obtained from a panel of various breast cancer cell lines (49) showed that TET2 CD cells activated genes encoding proteins enriched in the proteome of low tumorigenic cells and repressed genes encoding proteins that accumulate in highly tumorigenic cells (Supplementary Fig. 1C,D,E,F). Similar gene expression changes were also observed in decitabine-treated MCF-7 cells (Supplementary Fig. 1G). As a whole, this set of data indicates a lower aggressiveness of TET2 CD cells that correlates with a down-regulation of master regulators of cell growth and migration, likely contributing to their lower tumorigenic potential.

### TET2 CD triggers both activation and repression of CGI promoters

To test whether TET2 CD expression could reprogram CGIs, we next mapped 5hmCpGs genome-wide with a base-resolution method relying on selective chemical labeling (SCL) coupled to exonuclease digestion (SCL-exo, 19). Average profiles of SCL-exo signal centered on 28,691 hg19 CGIs showed an enrichment in 5hmCpGs specifically at CGIs from TET2 CD cells (Fig. 2A). Consistent with 5hmC occuring at methylated CpGs, average 5hmC enrichment was more pronounced at CGIs identified as methylated in MCF-7 cells (29; Fig. 2B). We then isolated 2,224 CGIs having at least 4 hydroxymethylated CpGs in TET2 CD cells. These 5hmC positive CGIs showed high SCL-exo signal (Fig. 2B) and 20% (448) of them were located within 500 bp of a transcription start site (TSS, Fig. 2C). However, these TSS-associated hydroxymethylated CGIs were found to associate poorly with highly regulated genes. Indeed, among the top 60 TET2 CD-regulated genes, only 5 up- and 2 down-regulated genes had a TSS associated with a 5hmC+ CGI (Fig. 2D). CGIs including TSSs, thereafter designated TSS-CGIs, were next classified into three groups (Fig. 2E and Supplementary Fig. 2A): (*i*) those having no detectable 5hmC in EV cells and gaining 5hmC in TET2 CD cells (low 5mC oxidation in EV cells, n=1034), (*ii*) those having readily detectable 5hmC in EV cells and losing 5hmC in TET2 CD cells (n=95), likely due to further oxidation of 5hmC by the expressed TET2 CD (moderate oxidation in EV cells), and (*iii*) those showing no signal at all in EV and TET2 CD cells (n=10,839) and which may correspond to CGIs subjected to a very high 5mC oxidation rate or being fully protected from methylation (no modification). Among low oxidation TSS-CGIs, only 34.5% were associated with a gene with detectable reads in RNA-seq, compared to 75% and 74% for the other two sets of TSS-CGIs, indicating that TET2-mediated gain in 5hmC at TSS-CGIs essentially targeted silent genes that did not get activated when demethylated. Accordingly, low oxidation TSS-CGIs tended to associate with the lowest expression quartiles of EV cell genes whereas moderate oxidation TSS-CGIs and unmodified TSS-CGIs were more associated with the highest expression quartiles (Fig. 2F). Fold-changes (TET2 CD vs EV) of genes associated with low and moderate oxidation TSS-CGIs were not significantly different (Fig. 2G), although their variance was (p < 0.0001). This was correlated to a higher representation of activated genes in low oxidation TSS-CGIs, whereas low and moderate oxidation TSS-CGIs equally associated with down-regulated genes (Fig. 2H). Altogether, these data indicate that 5mC oxidation dynamics at TSS-CGIs *per se* is not a robust predictor of the directionality of gene expression changes.

To further explore CGI chromatin remodeling upon TET2 CD expression, changes in the distribution of the active promoter mark H3K4me3 and the PRC2-mediated repression mark H3K27me3 were investigated by ChIP-seq in MCF-7 clones. Quite few genomic regions gained H3K4me3 (n=254) in TET2 CD cells compared to regions gaining H3K27me3 (n=1,939). However, 55.1% (140 out of 254) of these H3K4me3-up regions overlapped with CGIs compared to 11.3% (220 out of 1,939) for H3K27me3-up regions (Fig. 2I). Contrary to 5mC oxidation, changes in the levels of histone modifications reflected gene expression changes in TET2 CD cells *vs* EV cells, with CGIs gaining H3K4me3 being associated with activated genes and CGIs gaining H3K27me3 with repressed genes (Fig. 2J). In addition, genes gaining H3K4me3 in TET2 CD cells were not activated in TET2 mCD cells, whereas genes gaining H3K27me3 in TET2 CD cells tended to be repressed in TET2 mCD cells (Fig. 2J), suggesting that gene repression occurred also in the absence of 5mC oxidation, probably through protein/protein interaction between PRC2 and the catalytic domain of TET2. However, as verified by ChIP-qPCR on 2 down-regulated genes (*LHX2* and *JPH3*), the levels of both gene repression and H3K27 methylation were much lower in the absence of TET2 catalytic activity (Supplementary Fig. 2B,C,D). This was in accordance with previous studies suggesting that an active TET2 catalytic domain was required for deposition of H3K27me3 at CGIs (50). Interestingly, decitabine was not able to repress *LHX2* and *JPH3* (Supplementary Fig. 2E), indicating that TET2/PRC2 interaction in combination with DNA demethylation is probably required for repression of these genes. Consistent with these observations, mRNA levels of both *LHX2* and *JPH3* were significantly higher in breast cancer patient samples with low TET2 mRNA levels compared to high TET2 samples (Supplementary Fig. 2F).

As an example of TET2 CD-mediated gene activation, a subset of TSS-CGIs from the the breast cancer-methylated *PDCHA* locus (6) selectively gained H3K4me3 in TET2 CD cells in correlation with activation of the associated genes as shown for *PCDHA4* (Fig. 2K,L), *PCDHA3*, *PCDHA9* and *PCDHA10* (Fig. 2K, Supplementary Fig. 2E). Interrogation of TCGA RNA-seq data from breast cancer samples supported the data obtained in MCF-7 cells, showing that *PCDHA4* expression levels are positively correlated with *TET2* expression in patients (Fig. 2M). To unveil a potential dominant-negative function of TET2 mCD, we next focused on regions showing opposite H3K4me3 variations. Consistent with correlated gene repression and uncorrelated gene activation between TET2 CD and mCD cells (Fig. 2J), 73% (717 out of 983) of H3K4me3-down regions in TET2 mCD cells also lost H3K4me3 in TET2 CD cells, whereas a limited overlap was observed for H3K4me3-up regions (3.2%, 25 out of 764, Fig. 2N). Highlighting a potential dominant-negative function of TET2 mCD at a subset of sites, 19% (146 out of 764) of the H3K4me3-up regions in TET2 mCD compared to EV cells were called as H3K4me3-down in TET2 CD cells (Fig. 2N). Among the 136 annotated genes associated with these opposite H3K4me3 changes, 42 had both a TET2 mCD/TET2 CD fold change above 1.5 and a TET2 mCD/EV fold change above 1.2 (Fig. 2O). GO annotation for cellular components of these 42 genes with Pantherdb (25; http://pantherdb.org/) indicated a unique annotation for intracellular organelle (GO:0043229) with a 1.5 fold enrichment (FDR = 4.50e^-02^). In particular, from this list of 42 genes, 5 genes were involved in lysosome biogenesis and autophagy (*ATP6V0A2*, *LAMTOR4*, *PRKAB1*, *SLC15A4*, *SPPL3*, and *VPS33A*, Fig. 2P).

### TET2 CD alters 5mC oxidation and H3K4me3 levels at MYC binding sites

Since DNA methylation/demethylation can influence transcription factor binding to DNA (51, 52), high resolution SCL-exo data were next interrogated for enriched transcription factor binding sites (TFBSs) in TET2 CD cells. Iterative clustering of TET2 CD and EV 5hmCpGs using the heatmap clustering tool of Cistrome (30) isolated a set of CpGs (n = 24,008) with high 5hmC enrichment in TET2 CD cells compared to EV cells (Fig. 3A), and another set of CpGs (n = 18,404) with the opposite enrichment pattern and most likely corresponding to 5hmCpGs undergoing superoxidation in TET2 CD cells (Supplementary Fig. 3A). Analysis of these two populations of CpGs with TFmotifView (32) revealed a high enrichment (*p* = 0 for CpGs gaining 5hmC and *p* = 6.3e^-272^ for CpGs losing 5hmC) in the MYC binding E box CACGTG motif (Fig. 3B and Supplementary Fig. 3B). This is consistent with the observation that 72% of the MYC-bound TSSs are also engaged by TET2 in HEK293T cells (53). MYC ChIP-seq data (ENCODE, SRR575112.1) obtained from serum-fed MCF-7 cells were next used to generate a list of MYC-binding sites. These sites showed a higher oxidation of 5mC (as reflected by an increase in differential 5hmC levels) in TET2 CD cells compared to EV cells and a lower one in TET2 mCD cells, consistent with a dominant-negative effect of the catalytic dead mutant (Fig. 3C). CGIs were next categorized into MYC^high^ and MYC^low^ subsets based on MYC ChIP-seq data from MCF-7 cells (Supplementary Fig. 3C). MYC^high^ CGIs were depleted in 5hmC, whereas MYC^low^ sites were enriched in 5hmC (Fig. 3D and Supplementary Fig. 3C). A large fraction of MYC binding sites were associated with TSS-CGIs (Supplementary Fig. 3D) and, consistent with an inhibitory role of DNA methylation on MYC binding to DNA (51, 54), a higher engagement of MYC was found at TSS-CGIs with moderate 5mC oxidation compared to TSS-CGIs with low 5mC oxidation (Fig. 3E). Of note, as examplified in Fig. 3F, a large fraction of MYC-bound TSS-CGIs showed decreased levels of H3K4me3 in TET2 CD cells compared to EV cells, although other genomic regions showed mixed behaviors with CGIs either gaining H3K4me3 or showing no change (Supplementary Fig. 3E). Accordingly, MYC binding was detected at the center of H3K4me3-down regions and slightly off the center of H3K4me3-up regions, suggesting a strong relationship between promoter activity and MYC binding (Fig. 3G). TSSs from TSS-CGIs were next split into MYC^low^ and MYC^high^ subsets and analyzed for variation in H3K4me3 in TET2 CD and TET2 mCD cells compared to EV cells (Fig. 3H). Results showed that the degree of H3K4me3 loss at TSSs from TSS-CGIs in TET2 CD cells was correlated to the level of MYC binding in MCF-7 cells (Fig. 3I,J). Such a decrease in H3K4me3 levels was not observed in TET2 mCD cells, indicating a requirement for an active catalytic domain. As an example, the *SEZ6L2* and *KCTD13* TSS-CGIs, which both bind MYC in MCF-7 cells, lost H3K4me3 in TET2 CD cells compared to EV cells, whereas the *ASPHD1* TSS-CGI was not bound by MYC and gained H3K4me3 (Fig. 3K). Consistent with these observations, *SEZ6L2* and *KCTD13* showed a lower expression and *ASPHD1* a higher expression in TET2 CD cells (Fig. 3L). Collectively, these data are in strong support of an increased engagement of MYC at TSS-CGI binding sites upon 5mC oxidation by TET2 CD, leading to a lower transcription of the associated genes.

**Figure 3:**
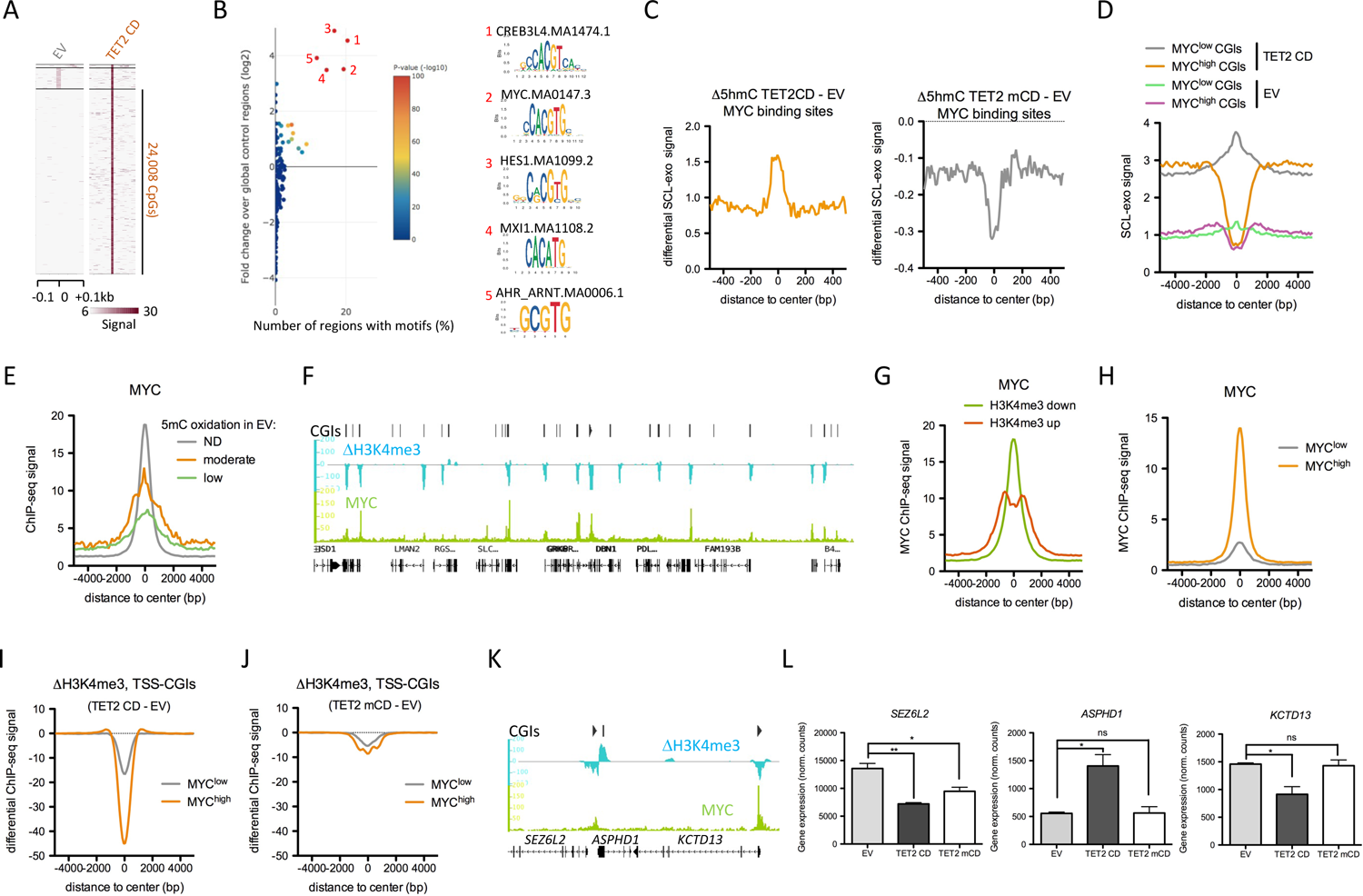
TET2 CD targets MYC binding sites. (**A**) Identification of CpGs gaining 5hmC in TET2 CD cells versus EV cells by heatmap clustering. (**B**) Enrichment of transcription factor binding motifs at CpGs gaining 5hmC in TET2 CD cells. Graph was generated with TFmotifView. (**C**) Differential SCL-exo signal between TET2 CD and EV cells (left panel) or TET2 mCD and EV cells (right panel) at MYC binding sites in MCF-7 cells (SRR575112.1). (**D**) SCL-exo signal at MYC^low^ and MYC^high^ CGIs in EV and TET2 CD cells. (**E**) Average MYC enrichment at CGIs classified according to their 5mC oxidation rate. (**F**) IGB representation of MYC ChIP-seq signal and differential H3K4me3 signal beteween TET2 CD and EV cells at a 400 kb region of chromosome 5. (**G**) Average MYC ChIP-seq signal at H3K4me3 up or down regions in TET2 CD cells compared to EV cells. (**H**) Average MYC ChIP-seq signal at MYC^high^ and MYC^low^ TSS-CGIs. (**I**,**J**) Differential H3K4me3 signal (**I**: TET2 CD - EV, **J**: TET2 mCD - EV) at MYC^high^ and MYC^low^ TSS-CGIs. (**K**) IGB snapshot of MYC ChIP-seq signal and H3K4me3 differential signal (TET2 CD - EV) at the *SEZ6L2*, *ASPHD1*, *KCTD13* locus. (**L**) *SEZ6L2*, *ASPHD1* and *KCTD13* mRNA levels in TET2 CD, mCD and EV cells (normalized RNA-seq read counts, mean +/- SEM, n=3).

### Decitabine and TET2 CD induce distinct cell reprogramming

Cancer alterations in DNA methylation can be counteracted by using DNA methyltransferase inhibitors like decitabine and 5-azacytidine (55). These drugs have been shown to promote tumor regression in hematological malignancies (56–58), and are under intensive investigation in solid tumors (59, 60). Combining HDAC inhibitors and DNA hypomethylating agents further reduced proliferation of lung cancer cancer cells by decreasing MYC levels and reversing immune evasion (60). Interestingly, DNMT inhibitors activate antiviral response genes through production of dsRNA from endogenous retroviral elements (ERVs), promoting apoptosis and immune checkpoint therapy in epithelial cancer cells (61–63). In order to compare the respective impact of TET2 CD expression and decitabine in MCF-7 cells, we first extracted DEGs (FC ≥ 2 and FC ≤ 0.5 respectively, adjusted *p* value ≤ 0.05) from public RNA-seq data of MCF-7 cells treated daily with 100 nM decitabine for 96 hours (48). Consistent with data obtained with other cell lines (48), the main outcome of decitabine in terms of gene regulation in MCF-7 cells was activation. Indeed, only 4.7% (80 out of 1704) of the DEGs were down-regulated by decitabine treatment (Fig. 4A). This was in striking contrast with TET2 CD DEGs which showed 58.5% (906 out of 1548) of down-regulated genes (Fig. 4A). In addition, the sets of DEGs poorly overlapped between decitabine treatment and TET2 CD with only 129 up-regulated genes in common, including *MCAM* (Fig. 4B,C). Conversely, *DSCR8*, a lncRNA known to activate WNT/*β*-catenin signaling in hepatocellular carcinoma (64), was confirmed to be massively induced by decitabine in MCF-7 cells by RT-qPCR (Fig. 4B,C). These data suggest that decitabine- and TET2 CD-induced transcriptional changes differ substantially.

**Figure 4:**
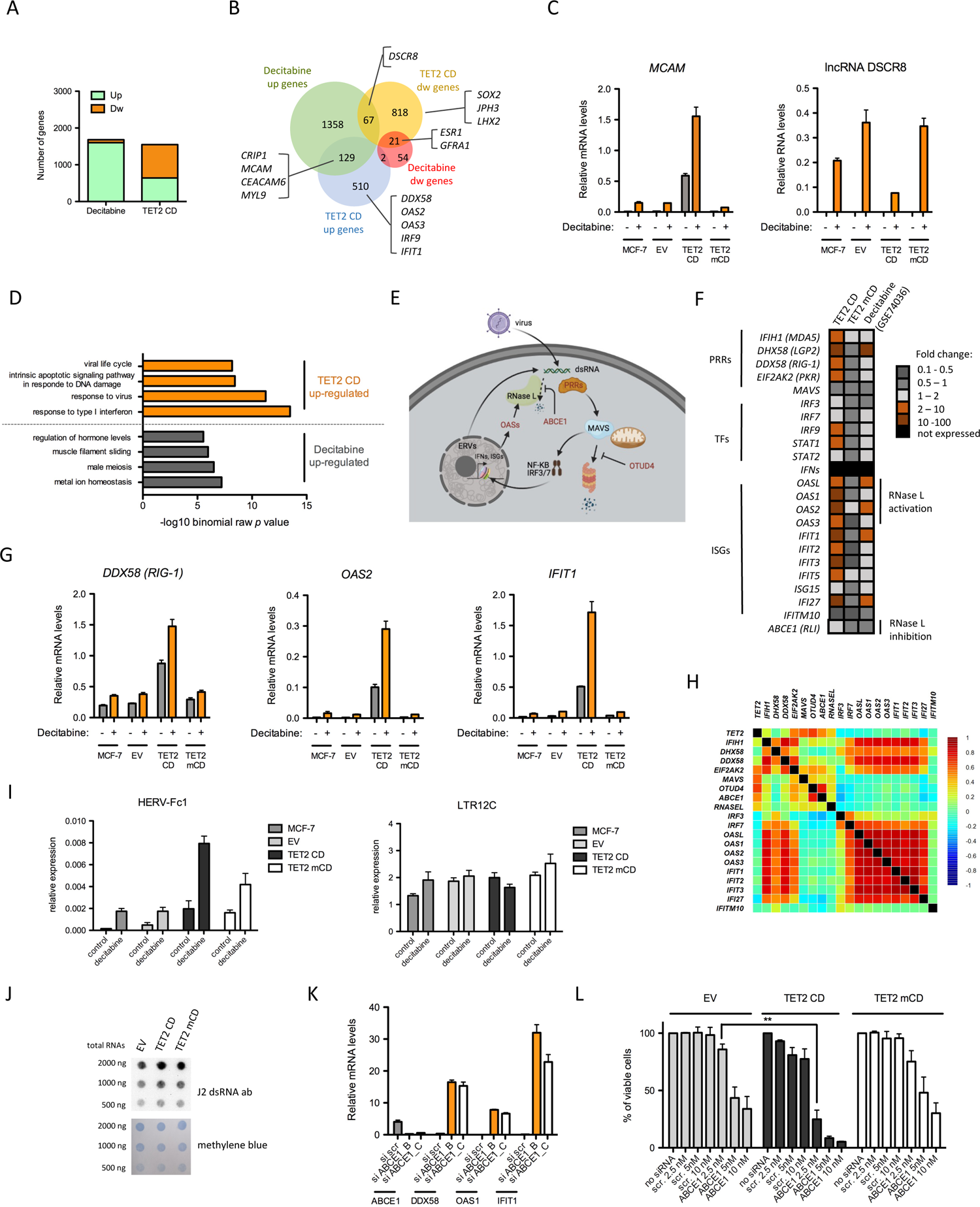
Differential transcriptional rewiring between TET2 CD expression and decitabine treatment. (**A**) Distribution of up- and down-regulated genes in DEGs from Decitabine-treated MCF-7 cells (GSE74036) and from TET2 CD cells *vs* EV cells. (**B**) Overlap between DEG lists from Decitabine-treated MCF-7 cells and from TET2 CD cells *vs* EV cells. A number of genes mentioned in the text are highlighted. Note that the *ESR1* gene encoding ER*α* is down-regulated both in Decitabine-treated MCF-7 cells and in TET2 CD cells. (**C**) RT-qPCR validation of the similar and opposite effects of decitabine and TET2 CD expression on *MCAM* and *DSCR8* RNA levels respectively (mean +/- SEM, n=3). (**D**) Functional annotation of TET2 CD and Decitabine up-regulated genes with GREAT. Only the top significantly enriched GO Biological processes are shown. (**E**) Outline of the antiviral reponse pathway. Double-stranded (ds) RNAs originating from viruses or from transcription of endogenous retroviral sequences (ERVs) can be sensed by pattern recognition receptors (PRRs) which activate the mitochondria-associated protein MAVS. Active MAVS is protected from degradation by OTUD4 and stimulates the nuclear translocation of IRF3, IRF7 and NF-KB transcription factors to induce expression of type I interferon genes (IFNs) and, in turn, interferon stimulated genes (ISGs). Among ISGs, OAS1,2,3 and L activate RNAse L that ultimately degrades dsRNAs. Activity of RNAse L can be counteracted by ABCE1 (figure made with BioRender). (**F**) Heatmap representation of the fold change of genes implicated in the type I interferon and antiviral response pathways. (**G**) RT-qPCR measurement of *DDX58*, *OAS2* and *IFIT1* in MCF-7 clones treated or not with 100 nM decitabine for 96 hours (mean +/- SEM, n=3). (**H**) Correlation heatmap between TET2 and genes from the antiviral response pathway in BRCA tumours (normal-like tumours, n=639). (**I**) RT-qPCR analysis of HERV-Fc1 and LTR12C endogenous retroviruses. Cells were treated with 100 nM decitabine for 96 hours (mean +/- SEM, n=3). (**J**) Dot blot analysis of dsRNA in total RNA from EV, TET2 CD and TET2 mCD cells. (**K**) RT-qPCR analysis of ABCE1, DDX58, OAS1 and IFIT1 in MCF-7 cells transfected either with a scrambled (scr) siRNA or with siRNA B and C targeting ABCE1 (mean +/- SEM, n=3). (**L**) Cell viability (MTT assay) after transfection of increasing concentrations of scrambled siRNA (scr) or ABCE1 siRNA C (mean +/- SEM, n=3).

Next, GO annotation of decitabine and TET2 CD induced genes revealed a TET2 CD-specific enrichment in antiviral response genes (Fig. 4D,E). Such genes included pattern recognition receptors (PRRs) involved in viral RNA sensing (*MDA5*, *LGP2*, *RIG-1* and *PKR*), transcription factors (*IRF9* and *STAT1*), and interferon stimulated genes (ISGs), whereas none of the interferon genes were induced (Fig. 4E,F), consistent with an already observed interferon-independent activation of ISGs (65). Of note, the four *OAS* genes which are involved in RNAse L activation through synthesis of 2’-5’-oligoadenylate (66) were highly induced in TET2 CD cells whereas the RNAse L inhibitor ABCE1 showed a moderate 2-fold increase (Fig. 4F). As already described in various cancer cell types, decitabine activated a subset of these genes (Fig. 4F). TET2 CD induction of *DDX58*, *OAS2* and *IFIT1* was confirmed by RT-qPCR, and these genes were further activated upon decitabine treatment (Fig. 4G). However, when looking at the correlation between TET2 expression and antiviral response genes in breast cancer patients, a positive correlation was found only for *EIF2AK2*, *MAVS*, *OTUD4*, *ABCE1* and *RNaseL* expression levels (Fig. 4H). In contrast, expression of ISGs in patients strongly correlated with expression of PRRs but not with expression of *MAVS*, *OTUD4*, *ABCE1* and *RNaseL*. One possible explanation could be that tumor cells with high PPRs, MAVS, OTUD4, ISGs and RNAseL are undergoing cell death and are thus counterselected. The anti-viral state triggered by decitabine has been shown to associate with an increased transcription of endogenous retroviruses (ERVs, 62,63). Consistent with these studies, transcription of HERV-Fc1 was increased by decitabine in our cell lines but basal expression was higher in TET2 CD cells as well as in TET2 mCD cells (Fig. 4I). Conversely, the LTR12C RNA levels were high in all conditions (Fig 4I). In agreement with these expression data, dot-blot analysis of dsRNA levels did not show dramatic differences between cell clones (Fig 4J). Considering that antiviral genes were not induced in TET2 mCD cells whereas HERV-Fc1 expression was increased, activation of the anti-viral state by TET2 CD may require additional mechanisms. Knowing that (i) viral and ERV RNAs are methylated in cells (67, 68), (ii) RNA methylation decreases the anti-viral response (69), and (iii) in ES cells TET2 oxidizes 5mC in ERV RNAs (68), it is then possible that the high anti-viral response in TET2 CD cells reflects a dual action of TET2 on both genomic DNA and RNA transcribed from repeated sequences. Such a scenario would be compatible with the additive effect observed when combining active TET2 CD and decitabine (Fig 4G,I). Of note, viral mimicry did not induce the death of TET2 CD cells, suggesting that the level of activation of the innate immune pathway remained below the threshold required for cell death commitment. Since RNAse L activation by 2’-5’-oligoadenylate is counteracted by the RNase L inhibitor RLI/ABCE1 (66, 70), we next tested the hypothesis that activation of the antiviral response pathway could sentitize TET2 CD cells to RNAse L-mediated cell death by transfecting siRNAs targeting ABCE1. Efficient knock-down of ABCE1 mRNA levels was observed, together with a massive induction of viral response genes in siRNA-transfected MCF-7 cells (Fig. 4K). MTT assays of cells challenged with increasing concentration of ABCE1 siRNAs next revealed an increased ability of ABCE1 knock-down to induce cell death in TET2 CD cells compared to EV and TET2 mCD cells (Fig. 4L). Collectively, these data indicate that enforced TET2 activity in MCF-7 breast cancer cells triggers a pre-activated antiviral state that predisposes cells to death induced by ABCE1 inactivation.

### TET2 regulates lysosome function

Innate immune response triggered by viral infection is associated with a RNase L-dependent autophagy of viral particles (71). Although autophagy was not a term enriched by gene ontology analysis of our RNA-seq data, a significant association of TET2 CD down-regulated genes (FC ≥ 2) with lysosome annotation was evidenced, and this association was even more pronounced when using a less stringent threshold of 1.5 fold decrease (Fig. 5A and Supplementary Fig. 4A). As already suggested in Fig. 2N,O,P, these data indicate that TET2 CD cells are endowed with altered lysosomal function. Down-regulation of *CLN3*, *CTSD* and *NAGLU* in TET2 CD cells was further confirmed by RT-qPCR analysis, and expression of these three genes was shown to be anti-correlated with TET2 expression levels in the breast cancer TCGA cohort of patients (n = 1,218), validating our *in vitro* observations (Fig. 5B,C and Supplementary Fig. 4B,C). Using the Pan-Cancer TCGA dataset gathering RNA-seq data from 11,060 patients, an anticorrelation between *TET2* expression and mRNA levels of lysosome proteins was confirmed for *CLN3*, *CTSD*, *CTSF*, *CTSZ*, *IFI30* and *NAGLU*, suggesting TET2 might down-regulate lysosomal genes in various types of cancer (Supplementary Fig. 4D). To interrogate a possible impact of TET2 CD expression on lysosome activity, acidic vesicles were next labeled with LysoTracker red (72). Data showed that acidic vesicle size was higher in TET2 CD cells, indicating a potential engorgement of lysosomes (Fig. 5D,E). Lysosomes often position next to the centrosome where they have a high probability to fuse with autophagosomes guided by molecular motors (73). Such a pericentrosomal positioning was obvious in EV, TET2 CD and TET2 mCD cells but a fraction of TET2 CD cells showed lysosomes that were scattered around the nucleus (Fig. 5D). Reduced levels of hydrolytic enzymes and mislocalized lysosomes have been observed in *CLN3* mutant cells (74) and engorged lysosomes were described in *SNX14* mutants causing cerebellar atrophy in human (75). Consistent with these observations, *SNX14* mRNA levels were also reduced in TET2 CD RNA-seq data (Fig. 5A).

**Figure 5:**
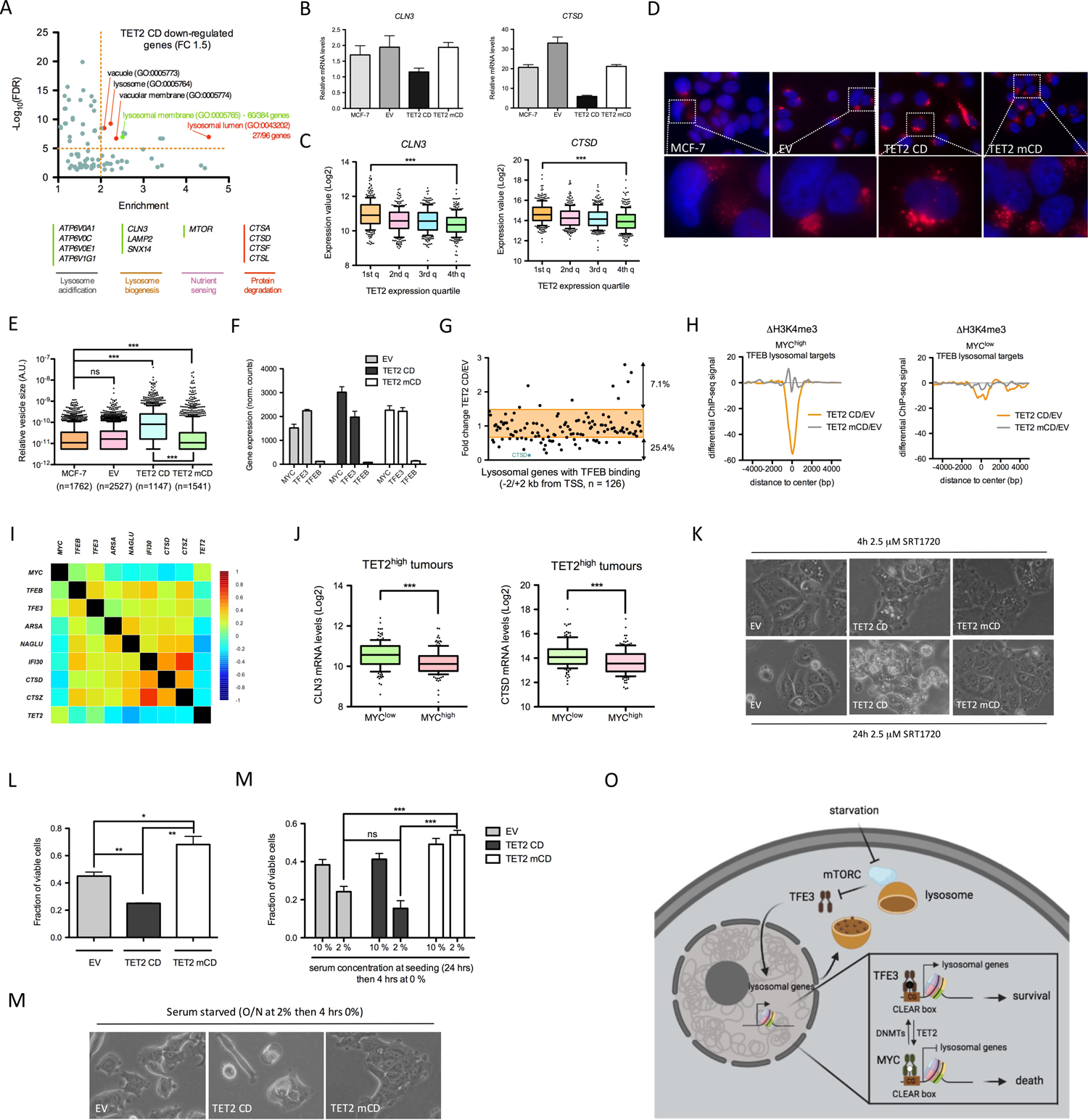
TET2 alters lysosome function. (**A**) GO cellular components annotation (Pantherdb) of 1.5 fold down-regulated genes in TET2 CD cells compared to EV cells. Specific down-regulated genes from the Lysosomal membrane and Lysosomal lumen annotations are shown as examples. (**B**) RT-qPCR analysis of CLN3 and CTSD mRNA levels in MCF-7 clones (mean +/- SEM, n=3). (**C**) Expression levels of *CLN3* and *CTSD* as a function of *TET2* mRNA levels ranked in quartiles (1st quartile: lowest expression, 4th quartile: highest expression) in BRCA tumours (TCGA BRCA dataset, n=1,218). (**D**) Lysotracker red labeling of acidic vesicles in MCF-7 clones. (**E**) Semi-quantification of acidic vesicle size in MCF-7 clones with ImageJ. (**F**) Expression levels (RNA-seq normalized read counts, mean +/- SEM, n=3) of transcription factors regulating lysosomal genes in MCF-7 clones. (**G**) Expression fold change in TET2 CD cells compared to EV cells of 126 genes associated with lysosome biogenesis and function and engaged by TFEB (ChIP-seq data from HUVECs) within 2 kb of their TSS. (**H**) Differential H3K4me3 ChIP-seq signal at MYC^high^ (left panel) and MYC^low^ (right panel) TSSs of TFEB lysosomal target genes. (**I**) Correlation heatmap between transcription factors regulating lysosome biogenesis and lysosomal genes in BRCA tumours (normal-like tumours, n=639). (**J**) CLN3 and CTSD mRNA levels in TET^high^ tumours (TCGA BRCA dataset, 4th quartile of expression, n=302) ranked according to MYC expression (n=151 for MYC^high^ and MYC^low^). (**K**) Representative images of MCF-7 clones treated for 4 hours or 24 hours with 2.5 μM SRT1720. (**L**) Cell viability (MTT assay) of the MCF-7 clones treated daily with 2.5 μM SRT1720 for 96 hours (mean +/- SEM, n=3). (**M**) Cell viability (MTT assay) of the MCF-7 clones grown for 24 hours in 10% or 2% serum and switched to serum-free medium for 4 hours (mean +/- SEM, n=3). (**N**) Representative images of MCF-7 clones grown for 24 hours in 2% serum and switched to serum-free medium for 4 hours. (**O**) Hypothetical model of the impact of TET2 on the coordinated transcription of lysosomal genes. Under starvation, TFE3 translocates to the nucleus where it activates lysosomal genes through binding to the CACGTG-containing CLEAR motif. Turnover of 5mC (black lollipop: 5mC, white lollipop: unmethylated C) at the CLEAR motif is controlled by the respective actions of DNMTs and TET2 and impacts the competitive binding of TFE3 and MYC, leading to an altered survival capability (figure made with BioRender).

Such an effect of TET2 CD expression on the activity of a large number of lysosomal genes was striking and suggestive of an alteration of a coordinated mechanism controlling lysosomal gene expression. It has been established that transcription factors from the basic helix-loop-helix leucine zipper (bHLH-ZIP) family such as TFEB and TFE3 coordinately activate lysosomal genes through binding of their basic domain to the CLEAR (Coordinated Lysosomal Expression and Regulation) motif (GTCACGTGAC) commonly found in the promoter of these genes (76, 77). In addition, an epigenetic mechanism involving MYC binding to the CLEAR motif (which contains the high affinity MYC binding site CACGTG) and recruitment of HDAC9 has been shown to antagonize the coordinated action of TFEB and TFE3 on lysosomal gene expression (78). Although MCF-7 cells did not express TFEB, they showed high levels of TFE3 and MYC mRNAs (Fig. 5F), suggesting that these two factors might compete for lysosomal gene regulation in these cells. TFEB ChIP-seq data obtained in HUVECs (GSM2354032) were then used to define a set of genes (n = 126) having TFEB binding sites within -/+2 kb from their TSS and belonging to the Lysosome gene set GO:0005764. Examination of these TFEB lysosomal targets revealed that 25.4% (32 out of 126) of them were down-regulated by expression of TET2 CD (Fig. 5G). Consistent with a role of MYC in shaping the chromatin landscape of these TFEB targets, MYC^high^ TSSs of TFEB lysosomal target genes showed a strong decrease in H3K4me3 levels in TET2 CD cells and a slight increase in TET2 mCD cells, whereas MYC^low^ TSS of TFEB lysosomal targets did not show variations in H3K4me3 levels (Fig. 5H). As expected, lysosomal gene mRNA levels were positively correlated with *TFEB* and *TFE3* levels and negatively correlated with MYC and TET2 levels in breast cancer patients (normal-like tumors, n = 639, Fig. 5I). In addition, TET^high^/MYC^high^ tumors had lower levels of *CLN3*, *CTSD* and *NAGLU* mRNAs compared to TET^high^/MYC^low^ tumors (Fig. 5J and Supplementary Fig. 4E), validating *in vivo* the hypothesis that TET2 repression of lysosomal genes is dependent on MYC.

We next challenged MCF-7 clones with SRT1720, a synthetic compound activating SIRT1 and known to activate autophagy and enhance lysosomal membrane permabilization, a process leading to the death of breast cancer cells (79). SRT1720 has been also shown to enhance TET2 enzymatic activity in myelodisplastic syndrome hematopoietic stem/progenitor cells (80). Thus, the impact of SRT1720 on global 5hmC levels in MCF-7 cells was first analyzed by dot blot. Data indicated that SRT1720, in MCF-7 cells, did not increase 5mC oxidation (Supplementary Fig. 4F), ruling out a possible regulation of TET activity by SRT1720 in these cells. At a concentration of 2.5 µM, SRT1720 induced the appearence of cytoplasmic vacuoles as soon as 4 hrs post-treatment and, whereas these vacuoles were cleared after 24 hrs in EV and TET2 mCD cells, they remained visible, together with hyper-vacuolized dead cells, in TET2 CD cells (Fig. 5K). After 4 days of treatment with daily doses of SRT1720, marked differences in cell survival were detected between clones, with a drastic reduction in viable TET2 CD cells compared to TET2 mCD cells (Fig. 5L). In the presence of SRT1720, addition of chloroquine, a drug that increases lysosome pH and inhibits fusion of autophagosomes with lysosomes (81), exacerbated the phenotype of TET2 CD cells which accumulated very large vacuoles (Supplementary Fig. 4G). In addition, serum starvation, a condition triggering autophagy through mTORC1 inhibition (82), induced high cell death rates in TET2 CD cells whereas TET2 mCD cells, likely through a dominant-negative function of the inactive catalytic domain, were protected from death (Fig. 5M,N). Collectively, these data indicated a prominent role of a TET2/MYC cross-talk in controlling lysosomal activity in breast cancer cells and impeding survival upon autophagy induction.

## DISCUSSION

*In vivo* DNA methylation dynamics relies in part on the respective levels of enzymes having opposite roles, namely DNMTs and TETs. Recent investigations using cell systems with combinatorial knock-out of these enzymes and live-cell imaging of DNA methylation reporters gave direct evidence for a cyclical behavior of DNA methylation with 5mC oxydation by TETs being a major contributor to the turnover of methylation at a genome-wide scale (83–86). CpGs that appear highly methylated at steady-state have low 5mC oxidation rate likely because they are poorly accessible to TETs. On the contrary, intermediate levels of methylation are reflecting a higher turnover thanks to the engagement of TETs. This is particularly true at enhancers which are major spots of 5mC oxidation in the genome (28,55,85). Promoter CGIs are sites of nucleosome depletion and as such should be highly accessible to TETs. Accordingly, the CXXC domain-containing TET1 and TET3 accumulate at TSS-CGIs (16, 87) where they are believed to protect DNA from aberrant methylation by the DNMTs. In addition, supported by the observed gain in DNA methylation upon TET2 knockout at a substantial number of CGIs (88), TET2 most likely also accumulate at TSS-CGIs. Although it does not contain a CXXC domain, TET2 could be targeted to TSS-CGIs through interaction with IDAX (89). Hence, promoter CGIs that are qualified as unmethylated in whole genome bisulfite sequencing experiments could be protected against DNA methylation through high 5mC oxidation rate. However, chromatin marks found at promoters (i.e. H3K4me3) can repress DNMT activity, providing an additional mechanism that could explain the lack of detectable DNA methylation at CGIs (90, 91). Here, by overexpressing either a catalytically active or inactive domain of TET2, we could highlight various operating modes of this protein. First, 5mC oxidation-dependent gene activation is observed at promoters that acquire H3K4 methylation with TET2 CD expression and not with TET2 mCD. Second, PRC2-associated gene repression (*i.e.* gain in H3K27me3) is partially 5mC oxidation-independent. Third, a number of gene repression events were mediated by an active 5mC oxidation mechanism as revealed by a decrease in H3K4me3 at TSS in TET2 CD cells and an opposite regulation in TET2 mCD cells. These antagonistic effects of TET2 CD versus mCD could reflect the recruitment of a transcriptional repressor that binds to unmethylated sequences in TET2 CD cells whereas demethylation would be impaired in TET2 mCD cells. We hypothesize that MYC could be one such factor since it is highly sensitive to DNA methylation (54), and engagement of MYC at TSSs induced either mild repression or activation (92). In favor of such a cross-talk between TET2 and MYC, we show that TET2 CD expression associates with demethylation of MYC binding motifs and coordinately represses genes involved in lysosome biogenesis and function, a characteristic that has also been assigned to MYC (78). Interestingly, MYC competes at the TSS of lysosomal genes with the activation factors TFEB and TFE3 (78). Since TFEB is not expressed in MCF-7 cells, the lysosomal transcriptional program is likely to be activated by TFE3 in these cells. Notably, TFE3 has been shown to bind both unmethylated and methylated CACGTG sites *in vitro* (57), although a negative impact of DNA methylation on TFE3 binding to DNA was also described (54). Based on these observations, we propose a model that positions the antagonistic effects of DNMTs and TETs at the CLEAR motif as a central regulatory switch for fine tuning of the lysosomal program (Fig. 5O). This switch would operate not only in breast cancer tumours or in other human tumour types, but also in other species for wich the lysosomal program is controlled through a competition between MYC and other bHLH-zip factors. In this regard, a remarkable enrichment of TET3 at TSSs of lysosomal genes (28% of the identified TET3 ChIP-seq peaks) in association with CACGTG motifs was described in mouse brain (87), reinforcing the hypothesis of a widespread involvement of TETs in controlling lysosomal functions.

The coordinated down-regulation of lysosomal genes by TET2 CD, although of a low magnitude for each individual gene, is likely to trigger a lysosomal storage disease-like state in breast cancer cells. *CTSD* appeared as the most affected gene in this process and is a central actor of lysosomal activity. *CTSD* knock-out mice develop a lysosomal storage disease that ultimately leads to death (93). In a mouse model of breast cancer, CTSD deficiency in the mammary epithelium impairs mTORC1 signaling and triggers the appearance of vacuolized cells with reduced proliferative activity upon serum starvation (94). TET2 CD cells were particularly prone to accumulate vacuoles, in particular when treated with the autophagy inducer SRT1720, and were highly sensitive to serum starvation. Cells respond to starvation by inhibiting lysosome-associated mTORC1, thus enhancing nuclear translocation of TFEB and TFE3 transcription factors and the activation of autophagy and lysosomal genes (95). Although an in depth characterization of the impact of TET2 CD expression on these complex pathways will be required to fully understand the phenotype of these cells, we propose that TET2 CD weakens lysosomal function and alters the cellular response to serum deprivation and autophagy induction. Our finding that TET2 expression, in the context of high MYC expression, negatively correlates with the mRNA levels of several lysosomal genes in breast tumours suggests that treatments combining autophagy inducers with DNA hypomethylating agents like decitabine and/or TET activating molecules could be beneficial to cancer patients. In this context, Vitamin C, a compound that has TET activating potential (96), could easily be administered to patients. Raising TET protein levels in tumours, through antimiR strategies against miRNAs targeting TET mRNAs (97), could provide an interesting alternative.

## DATA AVAILABILITY

All sequencing data are available at GEO (accession code GSE173344, https://www.ncbi.nlm.nih.gov/geo/). Unormalized wig files can be visualized at UCSC genome browser: https://genome.ucsc.edu/s/savner/MCF7_TET2 Flow cytometry data can be accessed at FlowRepository (https://flowrepository.org/) with the reference FR-FCM-Z446, reviewer access: https://flowrepository.org/id/RvFrDg4TjBSBZutxyIkLijvfWiuMcPcDaAiOmgI0wmqhoVhddiaSEbEbFDGxkFoF

## AKNOWLEDGEMENTS

We are grateful to Romain Gibeaux for his help with microscopy experiments. We thank Rémy Le Guével (ImPACcell, Biosit, Rennes) for his help in setting up cell migration assays, and members of the GEH (Rennes) and GenomEast Platform (IGBMC, Strasbourg) sequencing facilities.

## Supplementary Material

### Supplementary figure legends

**Supplementary Fig. 1:**
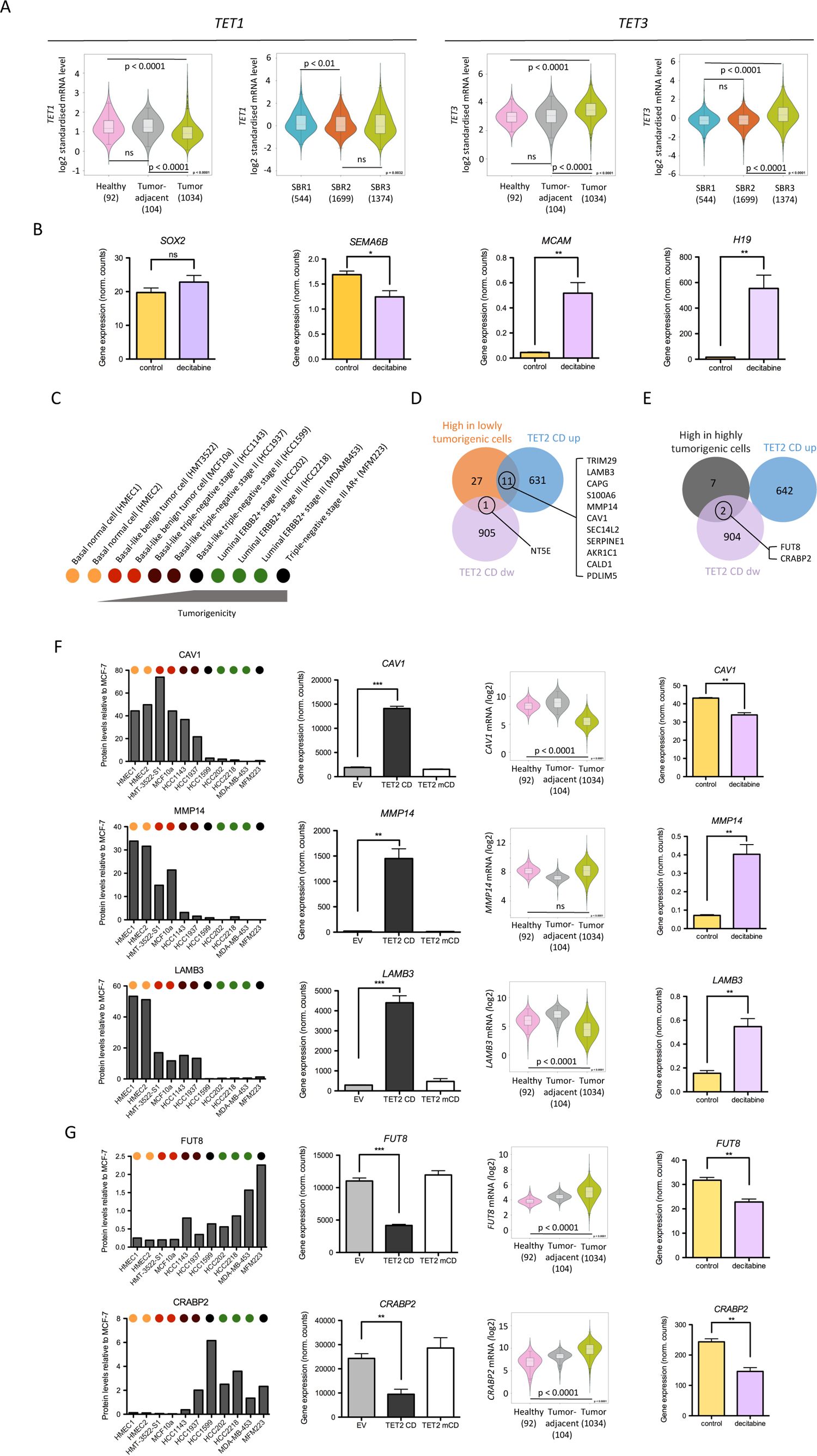
Ectopic expression of TET2 CD in MCF-7 cells correlates with lower tumorigenicity. **(A)** TET1 and TET3 expression in healthy tissue, tumor adjacent and tumors of BRCA patients as well as in BRCA tumors according to their Scarff Bloom and Richardson grade status. Violin plots were generated with Breast Cancer Gene-Expression Miner v4.5. (**C**) Diagram depicting the cell lines used by Geiger et al. (2012) to analyze the proteome of breast cancer cells and ranked according to their tumorigenicity. (**D**,**E**) Venn diagram showing the overlap between TET2 CD *vs* EV differentially expressed genes and proteins enriched in cell lines of low (**D**) or high (**E**) tumorigenicity. (**F**) Bar graph representation of selected protein levels (relative to their levels in MCF-7 cells), their corresponding mRNA levels in EV, TET2 CD and TET2 mCD cells (normalized RNA-seq read counts, mean +/- SEM, n=3), and violin plots of the levels of expression of their corresponding genes in healthy tissue, tumor adjacent and tumors of BRCA patients (protein levels in BRCA cell lines were recovered from Geiger et al., 2012). (**G**) Bar graph representation of selected mRNA levels upon decitabine treatment of MCF-7 cells (mean +/- SEM, n=3).

**Supplementary Fig. 2:**
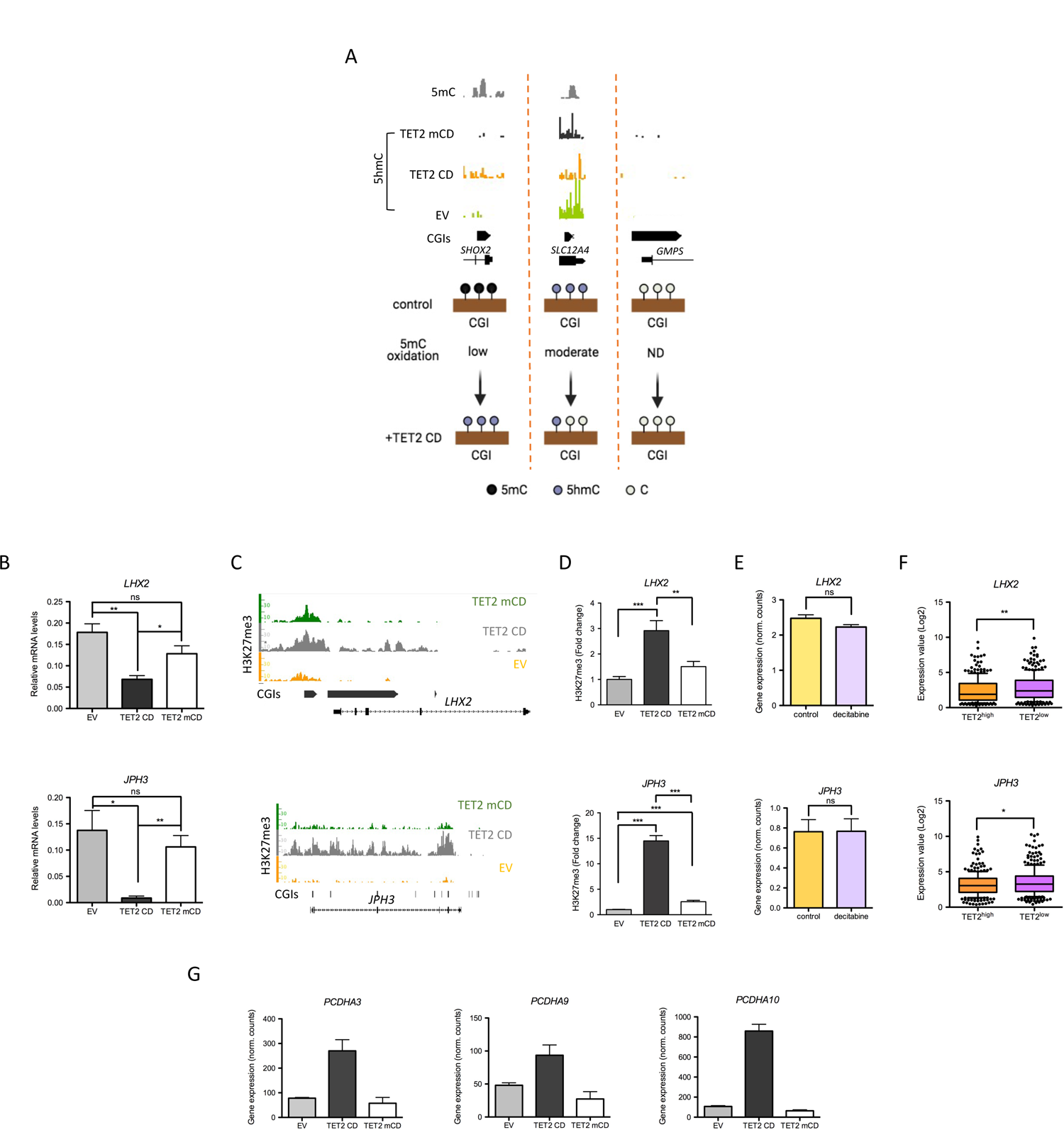
TET2 CD-induced chromatin changes at CGIs. (**A**) Classification of TSS-CGIs according to their 5mC oxidation rate. For each category, an example of MeDIP-seq (Ruike et al., 2010) and SCL-exo (5hmC) signals at one representative CGI is shown (IGB snapshot). (**B**,**C**,**D**) Expression levels (**B**, RT-qPCR of individual clones, mean +/- SEM, n=4 for EV and TET2 CD, n=3 for TET2 mCD), H3K27me3 ChIP-seq signal (**C,** IGB snapshots) and H3K27me3 ChIP-qPCR signal (**D,** mean +/- SEM, n=9) at 2 TET2 CD-repressed loci. (**E**) LHX2 and JPH3 mRNA levels in TCGA BRCA patient samples classified as TET^high^ (4th quartile of expression, n=300) or TET^low^ (1st quartiel of expression, n=300). (**F**) *PCDHA9* and *PCDHA10* mRNA levels in EV, TET2 CD and TET2 mCD cells (RNA-seq normalized read counts, mean +/- SEM, n=3).

**Supplementary Fig. 3:**
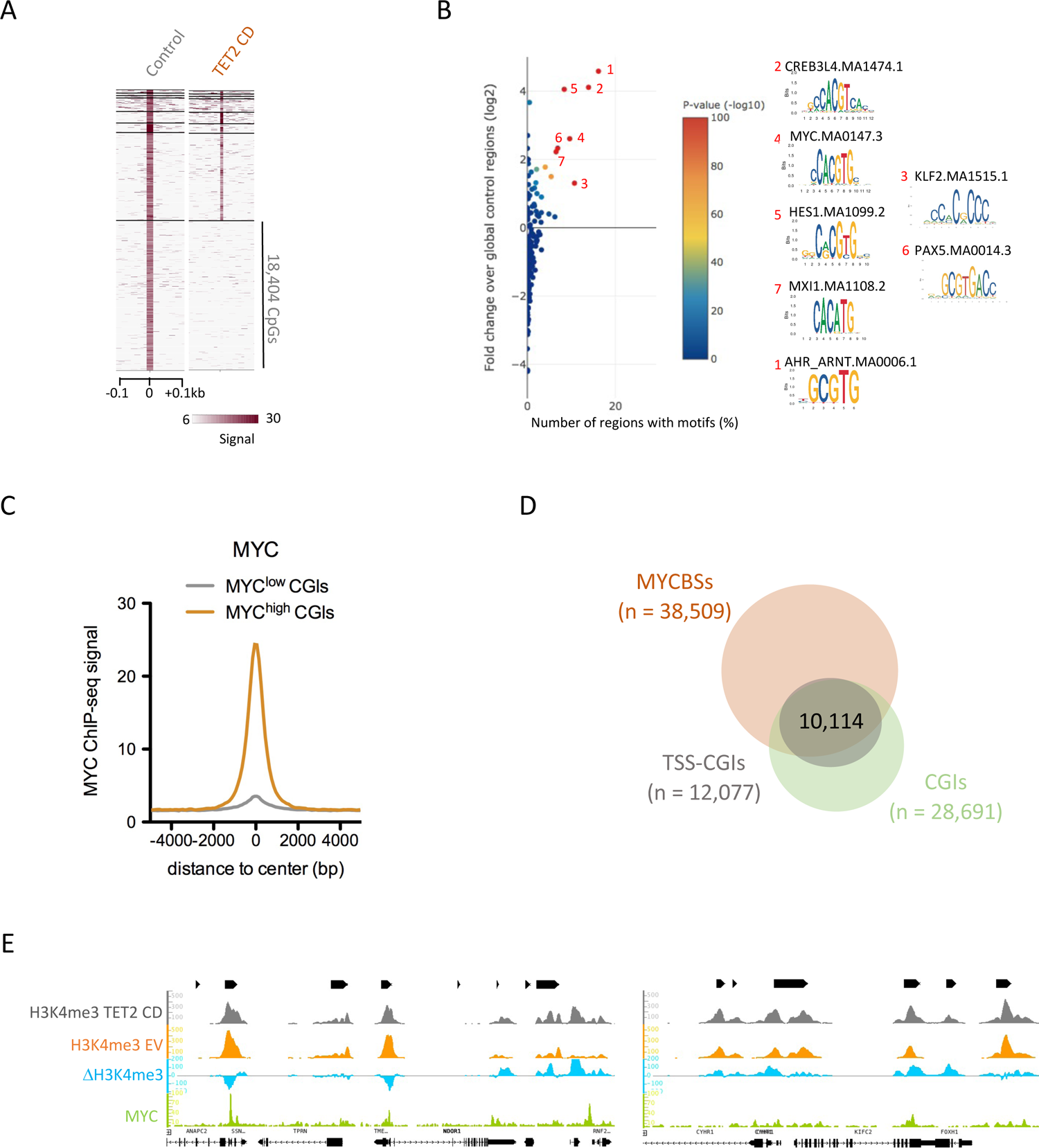
The MYC-binding E box motif (CACGTG) is enriched at TET2 CD targeted CpGs. (**A**) Identification of CpGs losing 5hmC in TET2 CD cells versus EV cells by heatmap clustering. (**B**) Enrichment of transcription factor binding motifs at CpGs losing 5hmC in TET2 CD cells. Graph was generated with TFmotifView. (**C**) Average MYC ChIP-seq signal at MYC^low^ and MYC^high^ CGIs. (**D**) Venn diagram showing the overlap between MYC binding sites (MYCBSs) and TSS-CGIs in MCF-7 cells. (**E**) IGB snapshots of H3K4me3 ChIP-seq signal in TET2 CD and EV cells and the corresponding differential signal (ΔH3K4me3, TET2 CD signal - EV signal), as well as MYC ChIP-seq signal in MCF-7 cells, at two different loci.

**Supplementary Fig. 4:**
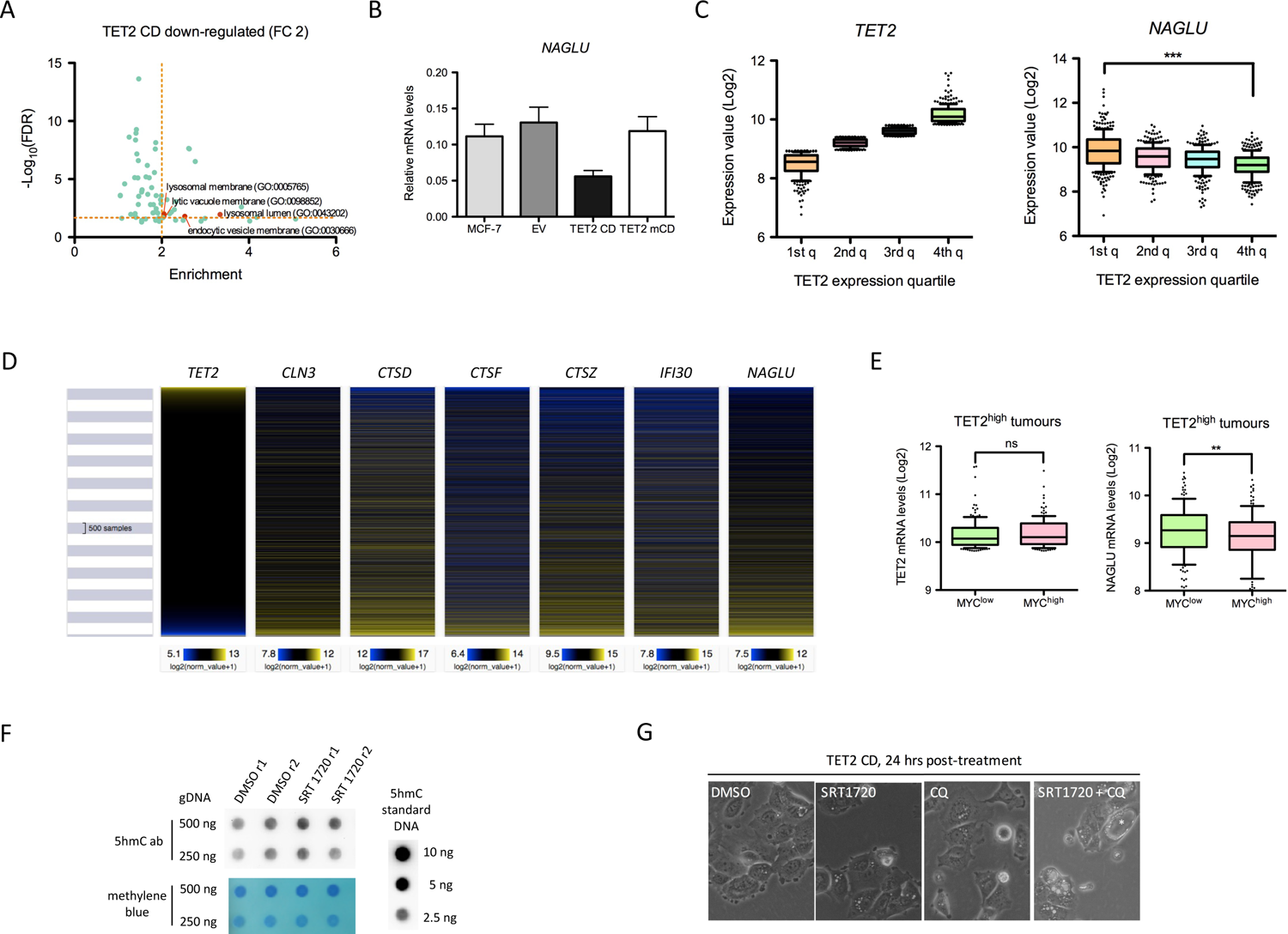
Ectopic expression of TET2 CD affects lysosomal function. (**A**) GO cellular components annotation (Pantherdb) of 2 fold down-regulated genes in TET2 CD cells compared to EV cells. (**B**) RT-qPCR analysis of NAGLU mRNA levels in MCF-7 clones (mean +/- SEM, n=3). (**C**) Expression levels of *TET2* and *NAGLU* as a function of *TET2* mRNA levels ranked in quartiles (1st quartile: lowest expression, 4th quartile: highest expression) in BRCA tumours (TCGA BRCA dataset, n=1,218). (**D**) Heatmap of *TET2* and selected lysosomal gene expression in TCGA pan-cancer samples (n=11,060). (**E**) TET2 and NAGLU mRNA levels in TET^high^ tumours (TCGA BRCA dataset, 4th quartile of expression, n=302) ranked according to MYC expression (n=151 for MYC^high^ and MYC^low^). (**F**) Dot blot analysis of 5hmC levels in MCF-7 cells treated with DMSO or 2.5 μM of SRT1720 for 48 hours. (**G**) Images of TET2 CD cells treated for 24 hours with DMSO, 2.5 μM SRT170, 10 μM chloroquine (CQ) or both. The white asterisk indicates a cell with a very large vacuole.

**Supplementary Table 1.** Top\_60 TET2 CD down-regulated genes

| geneSymbol | log2FoldChange | pval | adjPval | EV.rep1.norm | EV.rep2.norm | EV.rep3.norm | TET2_CD_rep1.norm | TET2_CD_rep2.norm | TET2_CD_rep3.norm | O/TS |
| --- | --- | --- | --- | --- | --- | --- | --- | --- | --- | --- |
| C1orf21 | -3,423897885 | 1,35E-132 | 1,24E-129 | 4158,566297 | 3729,427383 | 3631,806574 | 375,0255109 | 343,8892351 | 302,1171471 |  |
| USP2-AS1 | -5,368910617 | 1,42E-87 | 4,88E-85 | 548,6924619 | 496,3656432 | 512,0450067 | 12,69617615 | 7,316792237 | 3,492683781 |  |
| SEMA3D | -3,774436784 | 9,97E-86 | 3,27E-83 | 3380,990694 | 2748,39495 | 2407,388666 | 182,6296108 | 163,40836 | 213,0537107 | O |
| LINC00052 | -5,843969394 | 1,86E-75 | 4,84E-73 | 299,9518792 | 301,6632949 | 442,1029401 | 2,929886804 | 2,438930746 | 0 | O |
| TMEM229B | -2,198318873 | 1,66E-72 | 3,80E-70 | 584,2268309 | 645,943842 | 626,024671 | 135,7514219 | 130,8892833 | 126,6097871 |  |
| NOVA1 | -3,697266403 | 3,20E-70 | 7,11E-68 | 403,4196006 | 403,6104472 | 473,188303 | 33,20538378 | 29,26716895 | 23,57561552 | O |
| SLC27A6 | -2,522660009 | 8,46E-67 | 1,60E-64 | 655,2955688 | 523,9415122 | 493,048396 | 99,61615134 | 82,92364535 | 96,04880398 | O |
| KYNU | -2,65684218 | 1,13E-65 | 1,98E-63 | 15912,0814 | 13950,04722 | 12375,42839 | 2455,245142 | 2078,781972 | 1852,868746 |  |
| MUC15 | -5,097550132 | 2,84E-60 | 4,25E-58 | 234,1087838 | 280,772485 | 179,6043194 | 2,929886804 | 3,251907661 | 2,619512836 | O |
| ZMAT1 | -4,14872343 | 2,13E-58 | 3,03E-56 | 266,5077672 | 230,6345413 | 249,5463861 | 7,813031478 | 10,5686999 | 12,22439323 |  |
| DMKN | -2,479162795 | 3,68E-57 | 5,07E-55 | 1752,680493 | 1708,032617 | 1882,391424 | 398,4646054 | 280,4770357 | 238,3756681 | O |
| KAL1 | -2,636912012 | 6,04E-57 | 8,24E-55 | 606,1745294 | 563,2162348 | 583,714038 | 89,849862 | 85,3625761 | 90,80977831 | O |
| SOX2 | -6,939368588 | 4,10E-105 | 2,17E-102 | 742,0412342 | 649,2863716 | 786,6323797 | 0,976628935 | 3,251907661 | 3,492683781 | O |
| BEX4 | -2,913107922 | 6,31E-70 | 1,38E-67 | 916,5776936 | 849,0025142 | 731,3695122 | 106,4525539 | 119,5076065 | 87,31709453 | O |
| ARSK | -2,451038848 | 1,43E-69 | 3,08E-67 | 604,0842724 | 586,6139419 | 642,4308348 | 89,849862 | 115,442722 | 117,8780776 |  |
| HTR2C | -5,19333993 | 1,27E-66 | 2,36E-64 | 333,3959912 | 304,1701921 | 294,4474659 | 0,976628935 | 6,503815322 | 9,604880398 | O |
| RHOBTB3 | -2,604298213 | 5,63E-64 | 9,57E-62 | 46860,42651 | 51759,0706 | 57128,85277 | 6758,272228 | 8182,612652 | 9526,295013 | O |
| SGCG | -8,28613239 | 2,68E-181 | 9,22E-178 | 1857,193343 | 2028,079825 | 1879,800978 | 2,929886804 | 0,812976915 | 0,873170945 |  |
| GPC4 | -4,586971331 | 3,21E-166 | 6,33E-163 | 3516,857399 | 3419,407764 | 3160,345236 | 144,5410823 | 143,896914 | 92,5561202 | O |
| SLC6A14 | -6,008953689 | 1,90E-138 | 2,02E-135 | 6196,56687 | 6785,335055 | 6755,885553 | 82,03683052 | 73,98089928 | 71,60001751 | O |
| DSCAM-AS1 | -2,789784002 | 6,41E-126 | 5,20E-123 | 24762,22952 | 26065,04569 | 31464,4317 | 3970,973249 | 3577,911404 | 4027,0644 | O |
| GYG2 | -3,457269382 | 1,94E-125 | 1,49E-122 | 1261,470098 | 1087,993379 | 987,8237567 | 95,70963561 | 93,49234525 | 99,54148776 |  |
| GPER1 | -3,734482157 | 5,36E-113 | 3,70E-110 | 5785,83137 | 5634,669245 | 7122,865532 | 444,3661653 | 425,1869267 | 422,6147375 | O |
| RUNDC3B | -3,658346588 | 1,91E-112 | 1,25E-109 | 732,6350777 | 605,833487 | 646,7482463 | 42,97167313 | 57,72136098 | 44,53171821 |  |
| COLEC12 | -5,01543676 | 3,20E-111 | 2,01E-108 | 1683,702012 | 1604,4142 | 1278,817293 | 58,59773608 | 30,08014586 | 27,94147025 |  |
| ARMCX1 | -2,590631402 | 6,92E-110 | 3,97E-107 | 1503,93991 | 1504,973945 | 1671,701742 | 207,0453342 | 278,038105 | 269,8098221 | TS |
| STC1 | -4,161941245 | 4,91E-105 | 2,51E-102 | 4356,095583 | 5909,592304 | 5350,136361 | 288,1055357 | 237,3892592 | 247,1073775 | O |
| APLP1 | -3,294065236 | 1,57E-99 | 7,22E-97 | 1894,817969 | 2206,069525 | 2074,084496 | 208,998592 | 209,7480441 | 175,50736 |  |
| SEMA6B | -3,714511179 | 1,05E-98 | 4,69E-96 | 791,1622737 | 904,9898847 | 781,4514858 | 56,64447822 | 69,91601471 | 46,2780601 | O |
| STXBP1 | -2,151447001 | 4,42E-98 | 1,90E-95 | 2832,298232 | 2960,645579 | 2894,392686 | 611,3697132 | 655,2593937 | 656,6245509 |  |
| PCDH10 | -2,855712665 | 5,00E-96 | 2,03E-93 | 22099,24211 | 21945,37798 | 23426,27493 | 2417,156613 | 3161,667223 | 3381,791071 | TS |
| JPH3 | -4,889195383 | 1,87E-91 | 6,97E-89 | 609,3099149 | 646,7794744 | 575,9426973 | 16,60269189 | 16,2595383 | 13,97073512 | TS |
| SLC7A8 | -3,164737576 | 5,30E-89 | 1,92E-86 | 994,962331 | 1235,064681 | 1261,547647 | 140,6345666 | 116,2556989 | 111,765881 |  |
| KNDC1 | -3,518975828 | 5,91E-85 | 1,90E-82 | 515,24835 | 588,2852067 | 508,5910775 | 49,80807567 | 44,71373034 | 34,92683781 |  |
| MEIS3 | -3,540768756 | 1,39E-84 | 4,36E-82 | 1417,194245 | 1331,998039 | 1303,85828 | 126,9617615 | 100,8091375 | 90,80977831 |  |
| CRLF1 | -4,155436462 | 3,95E-84 | 1,21E-81 | 1280,282411 | 1496,617621 | 1358,257665 | 90,82649093 | 47,15266108 | 62,86830806 | O |
| PLXND1 | -1,890661743 | 2,15E-81 | 6,16E-79 | 19695,44656 | 21179,10307 | 19244,43013 | 5227,894688 | 4871,357676 | 5900,889248 | O |
| ABLIM1 | -2,801476743 | 2,00E-78 | 5,51E-76 | 1154,866991 | 1027,827847 | 1122,526996 | 125,9851326 | 139,0190525 | 186,8585823 |  |
| TNNT1 | -2,800243797 | 1,02E-74 | 2,60E-72 | 2358,855022 | 1951,201644 | 2055,951368 | 335,9603536 | 289,4197818 | 245,3610356 | O |
| SERPINA1 | -4,098577959 | 2,23E-74 | 5,50E-72 | 501,6616795 | 546,5035869 | 373,0243557 | 23,43909443 | 28,45419203 | 17,46341891 | O |
| DCLK1 | -4,582703693 | 9,28E-74 | 2,24E-71 | 2317,049882 | 1694,662499 | 1673,428707 | 83,01345945 | 56,90838406 | 47,15123105 | O |
| EVL | -2,945373313 | 7,51E-73 | 1,78E-70 | 17010,51145 | 16005,70291 | 17029,59802 | 2201,321619 | 2094,228534 | 1832,785814 | O |
| CA5B | -2,243132216 | 7,11E-72 | 1,61E-69 | 1661,754313 | 1597,729141 | 1663,066919 | 354,5163033 | 296,736574 | 363,2391132 |  |
| AMIGO2 | -3,723544536 | 1,75E-68 | 3,59E-66 | 8532,429065 | 9629,82773 | 7987,211319 | 509,8003039 | 614,6105479 | 609,4733198 | O |
| MECOM | -3,274252331 | 5,35E-68 | 1,07E-65 | 1020,045415 | 1003,594507 | 971,4175929 | 83,01345945 | 117,0686758 | 81,20489791 | O |
| CERS4 | -3,01214564 | 7,97E-68 | 1,57E-65 | 2505,173012 | 2799,368526 | 2286,501143 | 303,7315987 | 333,3205352 | 242,7415228 |  |
| EFNB3 | -3,133909387 | 1,19E-67 | 2,31E-65 | 802,6586872 | 774,631231 | 746,0487113 | 63,48088076 | 111,3778374 | 68,10733373 | O |
| MUC1 | -2,571423105 | 7,57E-67 | 1,45E-64 | 8563,78292 | 10625,06591 | 9083,833846 | 1377,046798 | 1424,335555 | 1758,566284 | O |
| PLXNA4 | -4,334818448 | 4,31E-66 | 7,92E-64 | 1393,156289 | 1334,504936 | 1196,786474 | 41,01841526 | 44,71373034 | 65,4878209 | O |
| SOX3 | -3,775269551 | 4,35E-64 | 7,50E-62 | 435,818584 | 387,7334317 | 547,4477812 | 30,27549698 | 18,69846905 | 34,92683781 | O |
| SCD5 | -3,061489362 | 1,08E-62 | 1,77E-60 | 1663,84457 | 1674,607321 | 1137,206195 | 138,6813087 | 187,7976674 | 168,5219924 | O |
| KIF1A | -3,695250631 | 2,18E-62 | 3,54E-60 | 2066,219042 | 2033,929252 | 2050,770474 | 177,7464661 | 134,9541679 | 100,4146587 |  |
| TBC1D9B | -1,002927084 | 3,20E-62 | 5,14E-60 | 13337,9299 | 14922,72333 | 13352,02688 | 6835,425914 | 7080,215954 | 6795,889467 |  |
| ARMCX4 | -4,971824518 | 1,31E-61 | 2,02E-59 | 339,6667621 | 461,2690825 | 563,853945 | 7,813031478 | 10,5686999 | 6,985367562 |  |
| GRIK2 | -4,464670505 | 2,17E-60 | 3,28E-58 | 357,4339466 | 310,0196189 | 343,6659573 | 9,766289347 | 11,38167681 | 12,22439323 | TS |
| CARNS1 | -3,279100185 | 4,37E-60 | 6,47E-58 | 229,9282698 | 223,9494821 | 253,8637976 | 24,41572337 | 19,51144596 | 21,82927363 |  |
| TATDN2P3 | -6,204977906 | 1,86E-57 | 2,59E-55 | 320,8544492 | 315,8690456 | 471,4613384 | 0 | 0 | 0 |  |
| CECR2 | -4,560948056 | 1,52E-56 | 2,03E-54 | 388,7878016 | 324,2253696 | 494,7753606 | 9,766289347 | 14,63358447 | 12,22439323 |  |
| CALCR | -3,737622625 | 5,57E-56 | 7,38E-54 | 1188,311103 | 1469,877384 | 1540,452432 | 70,3172833 | 92,67936833 | 99,54148776 | O |
| PDE9A | -4,201380737 | 1,05E-55 | 1,37E-53 | 419,0965281 | 396,925388 | 285,8126429 | 14,64943402 | 18,69846905 | 12,22439323 | O |

Top\_60 TET2 CD up-regulated genes
| geneSymbol | log2FoldChange | pval | adjPval | EV.rep1.norm | EV.rep2.norm | EV.rep3.norm | TET2_CD_rep1.norm | TET2_CD_rep2.norm | TET2_CD_rep3.norm | O/TS |
| --- | --- | --- | --- | --- | --- | --- | --- | --- | --- | --- |
| SCIN | 6,260110829 | 2,36E-259 | 3,25E-255 | 137,9569619 | 165,4552144 | 176,1503902 | 13925,75198 | 13978,32508 | 15457,74524 | O |
| SPANXA1 | 8,182277333 | 2,52E-212 | 1,16E-208 | 6,270770994 | 5,849426771 | 10,36178766 | 3603,760769 | 3519,377066 | 2841,298256 | TS |
| SPANXA2 | 8,182277333 | 2,52E-212 | 1,16E-208 | 6,270770994 | 5,849426771 | 10,36178766 | 3603,760769 | 3519,377066 | 2841,298256 |  |
| CAPN9 | 4,458695322 | 1,24E-179 | 3,41E-176 | 160,9497888 | 183,0034947 | 135,5667219 | 3224,828743 | 4341,296727 | 3871,639971 | TS |
| MROH2A | 6,695016996 | 1,85E-174 | 4,25E-171 | 25,08308397 | 20,8908099 | 16,40616379 | 3418,201272 | 2503,968899 | 2371,532287 |  |
| NPR1 | 4,88187124 | 2,93E-164 | 5,06E-161 | 43,89539695 | 46,79541417 | 40,58366833 | 1610,461113 | 1256,049334 | 1470,419872 |  |
| NFIA | 4,16191556 | 1,98E-158 | 3,03E-155 | 66,88822393 | 64,34369448 | 70,80554899 | 1291,103452 | 1230,034073 | 1321,10764 |  |
| FSTL1 | 4,68161344 | 3,89E-156 | 5,36E-153 | 55,39181044 | 44,28851698 | 34,53929219 | 1195,393816 | 1175,564619 | 1366,512529 | TS |
| CRIP1 | 6,334817782 | 8,55E-152 | 1,07E-148 | 238,2892978 | 238,1552328 | 213,2801293 | 19733,76426 | 23192,60544 | 30590,6709 | TS |
| KLHL7 | 2,254193146 | 1,10E-143 | 1,26E-140 | 332,3508627 | 346,7874443 | 318,6249705 | 1487,405868 | 1636,52253 | 1695,697976 |  |
| HOXC10 | 3,395680783 | 9,09E-136 | 8,95E-133 | 293,6811082 | 321,7184724 | 286,6761252 | 2795,112011 | 3675,468634 | 3464,742311 | O |
| MUC5AC | 6,651426964 | 1,47E-132 | 1,26E-129 | 50,16616795 | 64,34369448 | 37,12973911 | 7498,556961 | 7572,879965 | 8284,645929 |  |
| CAV1 | 2,837796847 | 4,29E-115 | 3,11E-112 | 1939,758494 | 2193,535039 | 1751,142114 | 15459,05941 | 14123,84795 | 13842,379 | TS and O |
| MMP14 | 5,737079984 | 1,48E-110 | 8,87E-108 | 11,49641349 | 26,74023667 | 25,04098684 | 1128,00642 | 1542,217208 | 1796,112634 | O |
| C4orf19 | 2,991300731 | 3,34E-105 | 1,84E-102 | 74,20412342 | 78,54944521 | 87,21171278 | 653,3647573 | 615,4235248 | 743,9416454 |  |
| LINC00857 | 5,191704197 | 9,32E-103 | 4,59E-100 | 10,45128499 | 10,02758875 | 9,498305353 | 426,7868445 | 530,8739256 | 454,0488916 | O |
| H19 | 5,585489819 | 2,37E-102 | 1,13E-99 | 281,1395662 | 391,9115937 | 258,1812091 | 21733,90031 | 26664,01686 | 11809,63704 | TS and O |
| CHST3 | 3,145688515 | 1,09E-97 | 4,55E-95 | 291,5908512 | 345,1161795 | 348,8468511 | 3251,197724 | 2668,190236 | 3270,02519 |  |
| C12orf56 | 5,34386373 | 5,90E-96 | 2,32E-93 | 8,361027991 | 2,506897188 | 8,634823048 | 404,324379 | 386,9770116 | 381,5757031 |  |
| PKP1 | 2,500452232 | 4,98E-92 | 1,91E-89 | 577,9560599 | 674,3553435 | 638,1134232 | 3762,951286 | 3477,102266 | 3754,635065 | TS |
| CYP26B1 | 4,512998315 | 1,38E-87 | 4,88E-85 | 25,08308397 | 29,24713386 | 35,4027745 | 782,2797767 | 747,938762 | 950,0099885 |  |
| GSTO2 | 1,655726411 | 2,48E-86 | 8,33E-84 | 546,6022049 | 500,5438051 | 524,9972413 | 1653,432787 | 1741,396552 | 1595,283317 |  |
| MYL9 | 7,406945569 | 4,81E-84 | 1,44E-81 | 0 | 0 | 0 | 443,3895364 | 432,5037189 | 462,780601 | O |
| ASCC1 | 1,890001573 | 7,95E-84 | 2,33E-81 | 1321,042423 | 1220,85893 | 1204,557815 | 4380,180772 | 5194,109511 | 4482,859633 |  |
| TRPM2 | 7,635985311 | 1,43E-79 | 4,02E-77 | 0 | 0 | 0 | 1355,560961 | 643,0647399 | 587,6440462 |  |
| CDCA7L | 3,09200055 | 1,15E-77 | 3,12E-75 | 363,7047176 | 344,2805471 | 357,4816742 | 2616,388916 | 3451,087005 | 3608,815517 | O |
| SCARA5 | 4,085520547 | 3,66E-76 | 9,71E-74 | 16,72205598 | 40,9459874 | 35,4027745 | 589,8838766 | 601,6029172 | 635,6684482 | TS |
| PARVB | 3,297086729 | 1,10E-74 | 2,75E-72 | 41,80513996 | 50,13794375 | 48,35500907 | 517,6133354 | 439,0075342 | 527,395251 | TS |
| PVT1 | 2,089502543 | 1,42E-72 | 3,31E-70 | 493,3006515 | 507,2288643 | 442,1029401 | 2294,101368 | 2003,988096 | 1971,619994 | TS |
| SPANXC | 6,128314802 | 2,61E-69 | 5,54E-67 | 5,225642495 | 6,685059167 | 3,453929219 | 839,9008839 | 686,1525164 | 440,0781564 | O |
| MEIS1 | 2,665208128 | 1,09E-68 | 2,28E-66 | 126,4605484 | 166,2908468 | 124,3414519 | 759,8173112 | 1003,213513 | 1014,624638 |  |
| CT45A2 | 6,831898267 | 2,33E-68 | 4,73E-66 | 0 | 0 | 0 | 258,8066677 | 269,0953589 | 317,8342241 |  |
| S100P | 3,721352249 | 8,48E-66 | 1,54E-63 | 488,075009 | 454,5840234 | 405,8366833 | 7396,010923 | 7304,597583 | 5684,342854 | O |
| COX20 | 1,51025022 | 9,78E-66 | 1,75E-63 | 546,6022049 | 599,9840602 | 553,4921574 | 1810,670045 | 1589,369869 | 1489,629633 |  |
| AC110619.1 | 2,85024139 | 1,07E-65 | 1,89E-63 | 174,5364593 | 174,6471707 | 173,5599433 | 1189,534043 | 1428,40044 | 1377,863752 |  |
| IFIT5 | 2,311082598 | 1,46E-63 | 2,45E-61 | 256,0564822 | 267,4023667 | 243,50201 | 1353,607704 | 1122,72112 | 1455,575966 | O |
| PEG10 | 4,77666762 | 1,48E-63 | 2,46E-61 | 106,6031069 | 105,2896819 | 146,7919918 | 3888,936418 | 4130,735706 | 5746,337991 | O |
| PARP10 | 4,568579349 | 4,14E-62 | 6,55E-60 | 48,07591095 | 49,30231136 | 62,17072595 | 1285,243678 | 1495,064547 | 2185,546876 | TS |
| SLC5A12 | 3,294019204 | 1,12E-61 | 1,75E-59 | 52,25642495 | 42,61725219 | 58,71679673 | 424,8335866 | 569,8968176 | 619,0782002 |  |
| C1orf116 | 5,048491193 | 1,29E-60 | 1,98E-58 | 33,44411197 | 35,09656063 | 34,53929219 | 2599,786224 | 1437,343186 | 904,6050993 |  |
| KLF4 | 2,619967108 | 1,39E-59 | 2,03E-57 | 965,698733 | 986,0462271 | 911,8373139 | 8001,520862 | 5374,590386 | 5145,596381 | TS and O |
| CIB2 | 4,342192095 | 3,25E-59 | 4,72E-57 | 14,63179898 | 11,69885354 | 15,54268149 | 396,5113475 | 379,6602194 | 281,1610444 | TS |
| CEACAM6 | 3,079913891 | 8,18E-59 | 1,17E-56 | 270,6882812 | 278,2655878 | 218,4610231 | 2318,517091 | 2914,522241 | 1842,390695 | O |
| ALDH3A1 | 5,845640638 | 1,59E-57 | 2,23E-55 | 4,180513996 | 7,520691563 | 2,590446914 | 733,44833 | 854,4387379 | 406,8976605 |  |
| SGCE | 2,938812139 | 7,26E-57 | 9,80E-55 | 58,52719594 | 53,48047334 | 35,4027745 | 374,048882 | 396,7327346 | 442,6976693 |  |
| SLC6A3 | 4,416265977 | 2,83E-53 | 3,46E-51 | 8,361027991 | 8,356323959 | 10,36178766 | 292,9886804 | 232,5113977 | 251,4732322 |  |
| PCDHB5 | 4,688584624 | 2,94E-53 | 3,56E-51 | 12,54154199 | 16,71264792 | 13,81571688 | 348,6565297 | 543,0685794 | 679,3269954 |  |
| AC110619.2 | 2,987306321 | 5,57E-53 | 6,62E-51 | 456,721154 | 431,1863163 | 436,9220462 | 3583,251562 | 4131,548683 | 3741,537501 |  |
| MSL3P1 | 1,61063399 | 5,97E-53 | 7,03E-51 | 786,9817597 | 766,274907 | 708,0554899 | 2735,537646 | 2126,74761 | 2131,410277 |  |
| IFI27 | 5,091839225 | 6,76E-53 | 7,83E-51 | 21,94769848 | 13,37011833 | 20,72357532 | 788,1395503 | 612,9845941 | 1650,293087 | O |
| MRPL23-AS1 | 4,942440792 | 6,94E-53 | 7,97E-51 | 9,40615649 | 5,849426771 | 6,044376134 | 275,4093596 | 393,480827 | 291,6390957 |  |
| MCAM | 4,311540974 | 2,46E-52 | 2,74E-50 | 24,03795548 | 25,90460427 | 23,31402223 | 488,3144674 | 543,8815563 | 877,5368 | O |
| SLC12A4 | 2,483549931 | 3,03E-52 | 3,34E-50 | 364,7498461 | 354,3081359 | 268,5429968 | 1755,002196 | 1922,690404 | 2115,6932 |  |
| VSIG2 | 5,559625875 | 6,12E-51 | 6,49E-49 | 8,361027991 | 2,506897188 | 0,863482305 | 301,7783408 | 388,6029655 | 447,063524 |  |
| GCNT1 | 1,816074645 | 9,30E-51 | 9,71E-49 | 484,9396235 | 508,0644967 | 494,7753606 | 1621,204032 | 1825,946152 | 1892,161438 | O |
| LAMB3 | 3,668614928 | 9,53E-50 | 9,59E-48 | 299,9518792 | 319,2115752 | 276,3143375 | 5208,362109 | 4373,815804 | 3977,293656 |  |
| CT45A1 | 6,102182684 | 2,07E-49 | 2,07E-47 | 0 | 0 | 0 | 194,349158 | 134,141191 | 211,3073688 | O |
| STS | 1,829549239 | 2,51E-49 | 2,49E-47 | 649,0247978 | 631,7380913 | 604,4376134 | 2584,160161 | 2270,644524 | 1983,844388 |  |
| CTB-167G5.5 | 3,495906374 | 4,68E-49 | 4,57E-47 | 81,52002292 | 85,23450438 | 47,49152676 | 632,8555497 | 1169,873781 | 1018,117322 |  |
| OLFML3 | 2,944408515 | 1,82E-48 | 1,77E-46 | 35,53436896 | 30,08276625 | 26,76795145 | 229,5077997 | 267,4694051 | 274,1756768 | O |

## REFERENCES

1. Cantone, I., and Fisher, A.G. (2013). Epigenetic programming and reprogramming during development. Nat. Struct. Mol. Biol. 20, 282–289.

2. Kulis, M., Heath, S., Bibikova, M., Queirós, A.C., Navarro, A., Clot, G., Martínez-Trillos, A., Castellano, G., Brun-Heath, I., Pinyol, M., et al. (2012). Epigenomic analysis detects widespread gene-body DNA hypomethylation in chronic lymphocytic leukemia. Nat. Genet. 44, 1236–1242.

3. Sánchez-Vega, F., Gotea, V., Margolin, G., and Elnitski, L. (2015). Pan-cancer stratification of solid human epithelial tumors and cancer cell lines reveals commonalities and tissue-specific features of the CpG island methylator phenotype. Epigenetics Chromatin 8, 14.

4. Bormann, F., Rodríguez-Paredes, M., Lasitschka, F., Edelmann, D., Musch, T., Benner, A., Bergman, Y., Dieter, S.M., Ball, C.R., Glimm, H., et al. (2018). Cell-of-origin DNA methylation signatures are maintained during colorectal carcinogenesis. Cell Rep. 23, 3407–3418.

5. Issa, J.P. (2004). CpG island methylator phenotype in cancer. Nat. Rev. Cancer 4, 988–993.

6. Novak, P., Jensen, T., Oshiro, M.M., Watts, G.S., Kim, C.J., and Futscher, B.W. (2008). Agglomerative epigenetic aberrations are a common event in human breast cancer. Cancer Res. 68, 8616–8625.

7. Dallosso, A.R., Hancock A.L., Szemes, M., Moorwood, K., Chilukamarri, L., Tsai, H.H., Sarkar, A., Barasch, J., Vuononvirta, R., Jones, C., et al. (2009). Frequent long-range epigenetic silencing of protocadherin gene clusters on chromosome 5q31 in Wilms’ tumor. PLoS Genet. 5(11):e10000745.

8. Wang, K.H., Lin, C.J., Liu, C.J., Liu, D.W., Huang, R.L., Ding D.C., Weng, C.F., and Chu, T.Y. (2015). Global methylation silencing of clustered proto-cadherin genes in cervical cancer: serving as diagnostic markers comparable to HPV. Cancer Med. 4, 43–55.

9. Vega-Benedetti, A.F., Loi, E., Moi, L., Blois, S., Fadda, A., Antonelli, M., Arcella, A., Badiali, M., Giangaspero, F., Morra, I., et al. (2019). Clustered protocadherins methylation alterations in cancer. Clin. Epigenetics 11, 100.

10. Fang, F., Turcan, S., Rimner, A., Kaufman, A., Giri, D., Morris, L.G., Shen, R., Seshan, V., Mo, Q., Heguy, A., et al. (2011). Breast cancer methylomes establish an epigenetic foundation for metastasis. Sci. Transl. Med. 3, 75ra25.

11. Moarii, M., Reyal, F., and Vert, J.P. (2015). Integrative DNA methylation and gene expression analysis to assess the universality of the CpG island methylator phenotype. Hum. Genomics 9, 26.

12. Huang, Y., and Rao, A. (2014). Connections between TET proteins and aberrant DNA modifications in cancer. Trends Genet. 30, 464–474.

13. Figueroa, M.E., Abdel-Wahab, O., Lu, C., Ward, P.S., Patel, J., Shih, A., Li, Y., Bhagwat, N., Vasanthakumar, A., Fernandez, H.F., et al. (2010). Leukemic IDH1 and IDH2 mutations result in a hypermethylation phenotype, disrupt TET2 function, and impair hematopoietic differentiation. Cancer Cell 18, 553–567.

14. Ichimura, N., Shinjo, K., An, B., Shimizu, Y., Yamao, K., Ohka, F., Katsushima, K., Hatanaka, A., Tojo, M., Yamamoto, E., et al. (2015). Aberrant TET1 methylation closely associated with CpG island methylator phenotype in colorectal cancer. Cancer Prev. Res. (Phila.) 8, 702–711.

15. Wu, X., and Zhang, Y. (2017). TET-mediated active DNA demethylation: mechanism, function and beyond. Nat. Rev. Genet. 18, 517–534.

16. Wu, H., D’Alessio, A.C., Ito, S., Xia, K., Wang, Z., Cui, K., Zhao, K., Sun, Y.E., and Zhang, Y. (2011). Dual functions of Tet1 in transcriptional regulation in mouse embryonic stem cells. Nature 473, 389–394 (2011).

17. Comşa, Ş., Cîmpean, A.M., and Raica, M. (2015). The story of MCF-7 breast cancer cell line: 40 years of experience in research. Anticancer Res. 35, 3147–3154.

18. Ito, S., Shen, L., Dai, Q., Wu, S.C., Collins, L.B., Swenberg, J.A., He, C., and Zhang, Y. (2011). Tet proteins can convert 5-methylcytosine to 5-formylcytosine and 5-carboxylcytosine. Science 333, 1300–1303.

19. Sérandour, A.A., Avner, S., Mahé, E.A., Madigou, T., Guibert, S., Weber, M., and Salbert, G. (2016). Single-CpG resolution mapping of 5-hydroxymethylcytosine by chemical labeling and exonuclease digestion identifies evolutionarily unconserved CpGs as TET targets. Genome Biol. 17, 56.

20. Langmead, B., Trapnell, C., Pop, M., and Salzberg, S.L. (2009). Ultrafast and memory-efficient alignment of short DNA sequences to the human genome. Genome Biol. 10, R25.

21. Li, B., and Dewey, C.N. (2011). RSEM: accurate transcript quantification from RNA-Seq data with or without a reference genome. BMC Bioinformatics 12, 323.

22. Love, M.I., Huber, W., and Anders, S. (2014) Moderated estimation of fold change and dispersion for RNA-seq data with DESeq2. Genome Biol. 15, 550.

23. Rau, A., Gallopin, M., Celeux, G., and Jaffrézic, F. (2013). Data-based filtering for replicated high-throughput transcriptome sequencing experiments. Bioinformatics 29, 2146–2152.

24. McLean, C.Y., Bristor, D., Hiller, M., Clarke, S.L., Schaar, B.T., Lowe, C.B., Wenger, A.M., and Bejerano, G. (2010). GREAT improves functional interpretation of cis-regulatory regions. Nat. Biotechnol. 28, 495–501.

25. Mi, H., Muruganujan, A., Casagrande, J.T., and Thomas, P.D. (2013). Large-scale gene function analysis with the PANTHER classification system. Nat. Protoc. 8, 1551–1566.

26. Li, H., Handsaker, B., Wysoker, A., Fennell, T., Ruan, J., Homer, N., Marth, G., Abecasis, G., Durbin, R., and 1000 Genome Project Data Processing Subgroup. (2009). The Sequence Alignment/Map format and SAMtools. Bioinformatics 25, 2078-2079.

27. Zhang, Y., Liu, T., Meyer, C.A., Eeckhoute, J., Johnson, D.S., Bernstein, B.E., Nusbaum, C., Myers, R.M., Brown, M., Li, W., and Liu, X.S. (2008). Model-based analysis of ChIP-Seq (MACS). Genome Biol. 9, R137.

28. Sérandour, A.A., Avner, S., Oger, F., Bizot, M., Percevault, F., Lucchetti-Miganeh, C., Palierne, G., Gheeraert, C., Barloy-Hubler, F., Péron, C.L., et al. (2012). Dynamic hydroxymethylation of deoxyribonucleic acid marks differentiation-associated enhancers. Nucleic Acids Res. 40, 8255–8265.

29. Ruike, Y., Imanaka, Y., Sato, F., Shimizu, K., and Tsujimoto, G. (2010). Genome-wide analysis of aberrant methylation in human breast cancer cells using methyl-DNA immunoprecipitation combined with high-throughput sequencing. BMC Genomics. 11, 137.

30. Liu, T., Ortiz, J.A., Taing, L., Meyer, C.A., Lee, B., Zhang, Y., Shin, H., Wong, S.S., Ma, J., Lei, Y., et al. (2011). Cistrome: an integrative platform for transcriptional regulation studies. Genome Biol. 12, R83.

31. Sérandour, A.A., Avner, S., and Salbert, G. (2018). Coupling exonuclease digestion with selective chemical labeling for base-resolution mapping of 5-hydroxymethylcytosine in genomic DNA. Bio-protocol 8, e2747.

32. Leporcq, C., Spill, Y., Balaramane, D., Toussaint, C., Weber, M., and Bardet, A.F. (2020). TFmotifView: a webserver for the visualization of transcription factor motifs in genomic regions. Nucleic Acids Res. 48, W208–W217.

33. Jézéquel, P., Campone, M., Gouraud, W., Guérin-Charbonnel, C., Leux, C., Ricolleau, G., and Campion, L. (2012). bc-GenExMiner: an easy-to-use online platform for gene prognostic analyses in breast cancer. Breast Cancer Res. Treat. 131, 765–775.

34. Huang, S., Zhu, Z., Wang, Y., Wang, Y., Xu, L., Chen, X., Xu, Q., Zhang, Q., Zhao, X., Yu, Y., and Wu, D. (2013). Tet1 is required for Rb phosphorylation during G1/S phase transition. Biochem. Biophys. Res. Commun. 434, 241–244.

35. Lian, C.G., Xu, Y., Ceol, C., Wu, F., Larson, A., Dresser, K., Xu, W., Tan, L., Hu, Y., Zhan, Q., et al. (2012). Loss of 5-hydroxymethylcytosine is an epigenetic hallmark of melanoma. Cell 150, 1135–1146.

36. Hsu, C.H., Peng, K.L., Kang, M.L., Chen, Y.R., Yang, Y.C., Tsai, C.H., Chu, C.S., Jeng, Y.M., Chen, Y.T., Lin, F.M., et al. (2012). TET1 suppresses cancer invasion by activating the tissue inhibitors of metalloproteinases. Cell Rep. 2, 568–579.

37. Wang, J., Sun, J., and Yang, F. (2020). The role of long non-coding RNA H19 in breast cancer. Oncol. Lett. 19, 7–16.

38. Miano, V., Ferrero, G., Reineri, S., Caizzi, L., Annaratone, L., Ricci, L., Cutrupi, S., Castellano, I., Cordero, F., and De Bortoli, M. (2016). Luminal long non-coding RNAs regulated by estrogen receptor alpha in a ligand-independent manner show functional roles in breast cancer. Oncotarget 7, 3201–3216.

39. Niknafs, Y.S., Han, S., Ma, T., Speers, C., Zhang, C., Wilder-Romans, K., Iyer, M.K., Pitchiaya, S., Malik, R., Hosono, Y., et al. (2016). The lncRNA landscape of breast cancer reveals a role for DSCAM-AS1 in breast cancer progression. Nat. Commun. 7, 12791.

40. Salameh, A., Fan, X., Choi, B.K., Zhang, S., Zhang, N., and An, Z. (2017). HER3 and LINC00052 interplay promotes tumor growth in breast cancer. Oncotarget 8, 6526–6539.

41. Ge, C., Li, Q., Wang, L., and Xu, X. (2013). The role of axon guidance factor semaphorin 6B in the invasion and metastasis of gastric cancer. J. Int. Med. Res. 41, 284–292.

42. Gurrapu, S., and Tamagnone, L. (2019). Semaphorins as regulators of phenotypic plasticity and functional reprogramming of cancer cells. Trends Mol. Med. 25, 303–314.

43. Toledano, S., Nir-Zvi, I., Engelman, R., Kessler, O., and Neufeld, G. (2019). Class-3 semaphorins and their receptors: potent multifunctional modulators of tumor progression. Int. J. Mol. Sci. 20, 556.

44. Mishra, R., Thorat, D., Soundararajan, G., Pradhan, S.J., Chakraborty, G., Lohite, K., Karnik, S., and Kundu, G.C. (2015). Semaphorin 3A upregulates FOXO 3a-dependent MelCAM expression leading to attenuation of breast tumor growth and angiogenesis. Oncogene 34, 1584–1595.

45. Dudzik, P., Trojan, S.E., Ostrowska, B., Lasota, M., Dulińska-Litewka, J., Laidler, P., and Kocemba-Pilarczyk, K.A. (2019). Aberrant promoter methylation may be responsible for the control of CD146 (MCAM) gene expression during breast cancer progression. Acta Biochim. Pol. 66, 619–625.

46. Stolzenburg, S., Rots, M.G., Beltran, A.S., Rivenbark, A.G., Yuan, X., Qian, H., Strahl, B.D., and Blancafort, P. (2012). Targeted silencing of the oncogenic transcription factor SOX2 in breast cancer. Nucleic Acids Res. 40, 6725–6740.

47. Li, X., Xu, Y., Chen, Y., Chen, S., Jia, X., Sun, T., Liu, Y., Li, X., Xiang, R., and Li, N. (2013). SOX2 promotes tumor metastasis by stimulating epithelial-to-mesenchymal transition via regulation of WNT/β-catenin signal network. Cancer Lett. 336, 379–389.

48. Sato, T., Cesaroni, M., Chung, W., Panjarian, S., Tran, A., Madzo, J., Okamoto, Y., Zhang, H., Chen, X., Jelinek, J., and Issa, J.J. (2017). Transcriptional selectivity of epigenetic therapy in cancer. Cancer Res. 77, 470–481.

49. Geiger, T., Madden, S.F., Gallagher, W.M., Cox, J., and Mann, M. (2012). Proteomic portrait of human breast cancer progression identifies novel prognostic markers. Cancer Res. 72, 2428–2439.

50. Kong, L., Tan, L., Lv, R., Shi, Z., Xiong, L., Wu, F., Rabidou, K., Smith, M., He, C., Zhang, L., et al. (2016). A primary role of TET proteins in establishment and maintenance of De Novo bivalency at CpG islands. Nucleic Acids Res. 44, 8682–8692.

51. Yin, Y., Morgunova, E., Jolma, A., Kaasinen, E., Sahu, B., Khund-Sayeed, S., Das, P.K., Kivioja, T., Dave, K., Zhong, F., et al. (2017). Impact of cytosine methylation on DNA binding specificities of human transcription factors. Science 356, eaaj2239.

52. Mahé, E.A., Madigou, T., and Salbert G. (2018). Reading cytosine modifications within chromatin. Transcription 9, 240–247.

53. Chen, L.-L., Lin, H.-P., Zhou, W.-J., He, C.-X., Zhang, Z.-Y., Cheng, Z.-L., Song, J.-B., Liu, P., Chen, X.-Y., Xia, Y.-K., et al. (2018). SNIP1 recruits TET2 to regulate c-MYC target genes and cellular damage response. Cell Rep. 25, 1485–1500.

54. Prendergast, G.C., and Ziff, E.B. (1991). Methylation-sensitive sequence-specific DNA binding by the c-Myc basic region. Science. 251, 186–189.

55. Jones, P.A., Issa, J.P., and Baylin, S. (2019). Targeting the cancer epigenome for therapy. Nat. Rev. Genet. 17, 630–641.

56. Von Hoff, D.D., Slavik, M., and Muggia, F.M. (1976). 5-Azacytidine. A new anticancer drug with effectiveness in acute myelogenous leukemia. Ann. Intern. Med. 85, 237–245.

57. Fenaux, P., Mufti, G.J., Hellstrom-Lindberg, E., Santini, V., Finelli, C., Giagounidis, A., Schoch, R., Gattermann, N., Sanz, G., List, A., et al. (2009). Efficacy of azacitidine compared with that of conventional care regimens in the treatment of higher-risk myelodysplastic syndromes: a randomised, open-label, phase III study. Lancet Oncol. 10, 223–232.

58. Issa, J.P., Garcia-Manero, G., Huang, X., Cortes, J., Ravandi, F., Jabbour, E., Borthakur, G., Brandt, M., Pierce, S., and Kantarjian, H.M. (2015). Results of phase 2 randomized study of low-dose decitabine with or without valproic acid in patients with myelodysplastic syndrome and acute myelogenous leukemia. Cancer 121, 556–561.

59. Nie, J., Liu, L., Li, X., and Han, W. (2014). Decitabine, a new star in epigenetic therapy: the clinical application and biological mechanism in solid tumors. Cancer Lett. 354, 12–20.

60. Topper, M.J., Vaz, M., Chiappinelli, K.B., DeStefano Shields, C.E., Niknafs, N., Yen, R.C., Wenzel, A., Hicks, J., Ballew, M., Stone, M., et al. (2017). Epigenetic therapy ties MYC depletion to reversing immune evasion and treating lung cancer. Cell 171, 1284–1300.e21.

61. Li, H., Chiappinelli, K.B., Guzzetta, A.A., Easwaran, H., Yen, R.W., Vatapalli, R., Topper, M.J., Luo, J., Connolly, R.M., Azad, N.S., et al. (2014). Immune regulation by low doses of the DNA methyltransferase inhibitor 5-azacitidine in common human epithelial cancers. Oncotarget 5, 587–598.

62. Chiappinelli, K.B., Strissel, P.L., Desrichard, A., Li, H., Henke, C., Akman, B., Hein, A., Rote, N.S., Cope, L.M., Snyder, A., et al. (2015). Inhibiting DNA methylation causes an interferon response in cancer via dsRNA including endogenous retroviruses. Cell 162, 974–986.

63. Roulois, D., Loo Yau, H., Singhania, R., Wang, Y., Danesh, A., Shen, S.Y., Han, H., Liang, G., Jones, P.A., et al. (2015). DNA-demethylating agents target colorectal cancer cells by inducing viral mimicry by endogenous transcripts. Cell 162, 961–973.

64. Wang, Y., Sun, L., Wang, L., Liu, Z., Li, Q., Yao, B., Wang, C., Chen, T., Tu, K., and Liu, Q. (2018). Long non-coding RNA DSCR8 acts as a molecular sponge for miR-485-5p to activate Wnt/β-catenin signal pathway in hepatocellular carcinoma. Cell Death Dis. 9, 851.

65. Yoneyama, M., Suhara, W., Fukuhara, Y., Fukuda, M., Nishida, E., and Fujita, T. (1998). Direct triggering of the type I interferon system by virus infection: activation of a transcription factor complex containing IRF-3 and CBP/p300. EMBO J. 17, 1087–1095.

66. Floyd-Smith, G., Slattery, E., and Lengyel, P. (1981). Interferon action: RNA cleavage pattern of a (2’-5’)oligoadenylate--dependent endonuclease. Science 212, 1030–1032.

67. Wnuk, M., Slipek, P., Dziedzic, M., and Lewinska, A. (2020). The Roles of Host 5-Methylcytosine RNA Methyltransferases during Viral Infections. Int. J. Mol. Sci. 21, 8176.

68. Guallar, D., Bi, X., Pardavila, J.A., Huang, X., Saenz, C., Shi, X., Zhou, H., Faiola, F., Ding, J., Haruehanroengra, P., et al. (2018). RNA-dependent chromatin targeting of TET2 for endogenous retrovirus control in pluripotent stem cells. Nat. Genet. 50, 443–451.

69. Stepanov, G., Zhuravlev, E., Shender, V., Nushtaeva, A., Balakhonova, E., Mozhaeva, E., Kasakin, M., Koval, V., Lomzov, A., Pavlyukov, M., et al. (2018). Nucleotide Modifications Decrease Innate Immune Response Induced by Synthetic Analogs of snRNAs and snoRNAs. Genes (Basel) 9, 531.

70. Le Roy, F., Bisbal, C., Silhol, M., Martinand, C., Lebleu, B., and Salehzada, T. (2001). The 2-5A/RNase L/RNase L inhibitor (RLI) pathway regulates mitochondrial mRNAs stability in interferon alpha-treated H9 cells. J. Biol. Chem. 276, 48473–48482.

71. Chakrabarti, A., Ghosh, P.K., Banerjee, S., Gaughan, C., and Silverman, R.H. (2012). RNase L triggers autophagy in response to viral infections. J. Virol. 86, 11311–11321.

72. Chazotte, B. (2011). Labeling lysosomes in live cells with LysoTracker. Cold Spring Harb. Protoc. pdb.prot5571.

73. Pu, J., Guardia, C.M., Keren-Kaplan, T., and Bonifacino, J.S. (2016). Mechanisms and functions of lysosome positioning. J. Cell Sci. 129, 4329–4339.

74. Schmidtke, C., Tiede, S., Thelen, M., Käkelä, R., Jabs, S., Makrypidi, G., Sylvester, M., Schweizer, M., Braren, I., Brocke-Ahmadinejad, N., et al. (2019). Lysosomal proteome analysis reveals that CLN3-defective cells have multiple enzyme deficiencies associated with changes in intracellular trafficking. J. Biol. Chem. 294, 9592–9604.

75. Akizu, N., Cantagrel, V., Zaki, M.S., Al-Gazali, L., Wang, X., Rosti, R.O., Dikoglu, E., Gelot, A.B., Rosti, B., Vaux, K.K., et al. (2015). Biallelic mutations in SNX14 cause a syndromic form of cerebellar atrophy and lysosome-autophagosome dysfunction. Nat. Genet. 47, 528–534.

76. Sardiello, M., Palmieri, M., di Ronza, A., Medina, D.L., Valenza, M., Gennarino, V.A., Di Malta, C., Donaudy, F., Embrione, V., Polishchuk, R.S., et al. (2009). A gene network regulating lysosomal biogenesis and function. Science 325, 473–477.

77. Perera, R.M., Stoykova, S., Nicolay, B.N., Ross, K.N., Fitamant, J., Boukhali, M., Lengrand, J., Deshpande, V., Selig, M.K., Ferrone, C.R., et al. (2015). Transcriptional control of autophagy-lysosome function drives pancreatic cancer metabolism. Nature 524, 361–365.

78. Annunziata, I., van de Vlekkert, D., Wolf, E., Finkelstein, D., Neale, G., Machado, E., Mosca, R., Campos, Y., Tillman, H., Roussel, M.F., et al. (2019). MYC competes with MiT/TFE in regulating lysosomal biogenesis and autophagy through an epigenetic rheostat. Nat. Commun. 10, 3623.

79. Lahusen, T.J., and Deng, C.X. (2014). SRT1720 induces lysosomal-dependent cell death of breast cancer cells. Mol. Cancer Ther. 14, 183–192.

80. Sun, J., He, X., Zhu, Y., Ding, Z., Dong, H., Feng, Y., Du, J., Wang, H., Wu, X., Zhang, L., et al. (2018). SIRT1 activation disrupts maintenance of myelodysplastic syndrome stem and progenitor cells by restoring TET2 function. Cell Stem Cell 23, 355–369.e9.

81. Mauthe, M., Orhon, I., Rocchi, C., Zhou, X., Luhr, M., Hijlkema, K.J., Coppes, R.P., Engedal, N., Mari, M., and Reggiori, F. (2018). Chloroquine inhibits autophagic flux by decreasing autophagosome-lysosome fusion. Autophagy 14, 1435–1455.

82. Yim, W.W., and Mizushima, N. (2020). Lysosome biology in autophagy. Cell Discov. 6, 6.

83. Rulands, S., Lee, H.J., Clark, S.J., Angermueller, C., Smallwood, S.A., Krueger, F., Mohammed, H., Dean, W., Nichols, J., Rugg-Gunn, P., et al. (2018). Genome-scale oscillations in DNA methylation during exit from pluripotency. Cell Syst. 7, 63–76.e12.

84. Song, Y., van den Berg, P.R., Markoulaki, S., Soldner, F., Dall’Agnese, A., Henninger, J.E., Drotar, J., Rosenau, N., Cohen, M.A., Young, R.A., et al. (2019). Dynamic enhancer DNA methylation as basis for transcriptional and cellular heterogeneity of ESCs. Mol. Cell. 75, 905–920.e6.

85. Ginno, P.A., Gaidatzis, D., Feldmann, A., Hoerner, L., Imanci, D., Burger, L., Zilbermann, F., Peters, A.H.F.M., Edenhofer, F., Smallwood, S.A., et al. (2020). A genome-scale map of DNA methylation turnover identifies site-specific dependencies of DNMT and TET activity. Nat. Commun. 11, 2680.

86. Charlton, J., Jung, E.J., Mattei, A.L., Bailly, N., Liao, J., Martin, E.J., Giesselmann, P., Brändl, B., Stamenova, E.K., Müller, F.J., et al. (2020). TETs compete with DNMT3 activity in pluripotent cells at thousands of methylated somatic enhancers. Nat. Genet. 52, 819–827.

87. Jin, S.G., Zhang, Z.M., Dunwell, T.L., Harter, M.R., Wu, X., Johnson, J., Li, Z., Liu, J., Szabó, P.E., Lu, Q., et al. (2016). Tet3 Reads 5-Carboxylcytosine through Its CXXC Domain and Is a Potential Guardian against Neurodegeneration. Cell Rep. 14, 493–505.

88. Zhang, Y.W., Wang, Z., Xie, W., Cai, Y., Xia, L., Easwaran, H., Luo, J., Yen, R.-W.C., Li, Y., and Baylin, S.B. (2017). Acetylation enhances TET2 function in protecting against abnormal methylation during oxidative stress. Mol. Cell 65, 323–335.

89. Ko, M., An, J., Bandukwala, H.S., Chavez, L., Äijö, T., Pastor, W.A., Segal, M.F., Li, H., Koh, K.P., Lähdesmäki, H., et al. (2013). Modulation of TET2 expression and 5-methylcytosine oxidation by the CXXC domain protein IDAX. Nature 497, 122–126.

90. Guo, X., Wang, L., Li, J., Ding, Z., Xiao, J., Yin, X., He, S., Shi, P., Dong, L., Li, G., et al. (2015). Structural insight into autoinhibition and histone H3-induced activation of DNMT3A. Nature 517, 640–644.

91. Masalmeh, R.H.A., Taglini, F., Rubio-Ramon, C., Musialik, K.I., Higham, J., Davidson-Smith, H., Kafetzopoulos, I., Pawlicka, K.P., Finan, H.M., Clark, R., et al. (2021). De novo DNA methyltransferase activity in colorectal cancer is directed towards H3K36me3 marked CpG islands. Nat Commun. 12, 694.

92. Walz, S., Lorenzin, F., Morton, J., Wiese, K.E., von Eyss, B., Herold, S., Rycak, L., Dumay-Odelot, H., Karim, S., Bartkuhn, M., et al. (2014). Activation and repression by oncogenic MYC shape tumour-specific gene expression profiles. Nature 511, 483–487.

93. Koike M, Nakanishi H, Saftig P, Ezaki J, Isahara K, Ohsawa Y, Schulz-Schaeffer W, Watanabe T, Waguri S, Kametaka S, et al. (2000). Cathepsin D deficiency induces lysosomal storage with ceroid lipofuscin in mouse CNS neurons. J. Neurosci. 20, 6898–6906.

94. Ketterer, S., Mitschke, J., Ketscher, A., Schlimpert, M., Reichardt, W., Baeuerle, N., Hess, M.E., Metzger, P., Boerries, M., Peters, C., et al. (2020). Cathepsin D deficiency in mammary epithelium transiently stalls breast cancer by interference with mTORC1 signaling. Nat Commun. 11, 5133.

95. Martina, J.A., Diab, H.I., Lishu, L., Jeong-A, L., Patange, S., Raben, N., and Puertollano, R. (2014). The nutrient-responsive transcription factor TFE3 promotes autophagy, lysosomal biogenesis, and clearance of cellular debris. Sci. Signal. 7, ra9.

96. Yue, X., and Rao, A. (2020). TET family dioxygenases and the TET activator vitamin C in immune responses and cancer. Blood 136, 1394–1401.

97. Fuentes-Mattei, E., Bayraktar, R., Manshouri, T., Silva, A.M., Ivan, C., Gulei, D., Fabris, L., Soares do Amaral, N., Mur, P., Perez, C., et al. (2020). miR-543 regulates the epigenetic landscape of myelofibrosis by targeting TET1 and TET2. JCI Insight 5:e121781.

